# Bacterial chemotaxis to saccharides is governed by a trade-off between sensing and uptake

**DOI:** 10.1101/2021.11.05.467490

**Authors:** Noele Norris, Uria Alcolombri, Johannes M. Keegstra, Yutaka Yawata, Filippo Menolascina, Emilio Frazzoli, Naomi M. Levine, Vicente I. Fernandez, Roman Stocker

## Abstract

To swim up gradients of nutrients, *E. coli* senses nutrient concentrations within its periplasm. For small nutrient molecules, periplasmic concentrations typically match extracellular concentrations. However, this is not necessarily the case for saccharides, such as maltose, which is transported into the periplasm via a specific porin. Previous observations have shown that under various conditions *E. coli* limits maltoporin abundance so that, for extracellular micromolar concentrations of maltose, there are predicted to be only nanomolar concentrations of free maltose in the periplasm. Thus, in the micromolar regime, the total uptake of maltose from the external environment into the cytoplasm is limited not by the abundance of cytoplasmic transport proteins but by the abundance of maltoporins. Here we present results from experiments and modeling showing that this porin-limited transport enables *E. coli* to sense micromolar gradients of maltose despite having a high-affinity ABC transport system that is saturated at these micromolar levels. We used microfluidic assays to study chemotaxis of *E. coli* in various gradients of maltose and methyl-aspartate and leveraged our experimental observations to develop a mechanistic transport-and-sensing chemotaxis model. Incorporating this model into agent-based simulations, we discover a trade-off between uptake and sensing: although high-affinity transport enables higher uptake rates at low nutrient concentrations, it severely limits dynamic sensing range. We thus propose that *E. coli* may limit periplasmic uptake to increase its chemotactic sensitivity, enabling it to use maltose as an environmental cue.

**Statement of Significance:** Bacterial chemotaxis is among the best-studied systems in biology and is paradigmatic of the mechanisms used by cells to link sensory inputs with regulated responses, thus providing insight into the ecological basis of cellular physiology. Here we present a mechanistic chemotaxis model that describes how the regulation of the transport of a sugar into and out of the cell’s periplasm affects the cell’s motile response to that sugar. Based on observations from population-level chemotaxis assays, we uncover an ecologically relevant trade-off between sensing and uptake. The general finding of this work is that, while high-affinity transport allows for higher uptake rates, it can severely limit the cell’s dynamic sensing range.

## Introduction

Many bacterial species can swim to actively seek environments favorable for growth. These species employ chemotaxis, in which they follow chemical gradients by biasing their swimming direction in response to temporal measurements of their environment (1). Chemotaxis allows cells to find and exploit chemicals in complex landscapes, such as the ocean or the human gut (2, 3). This makes them excellent microscale source-seekers – a trait that could allow bacteria to be re-engineered and deployed as “microbots” for a variety of tasks, such as bioremediation (4–6) and targeted medical treatment (7, 8). The chemicals that act as attractants for chemotaxis are often metabolic resources for bacteria. However, the precise relationship between chemotaxis and the benefit in confers, either directly through increased uptake of nutrients or indirectly as sensory cues that direct bacteria into more favorable environments, is less clear and has become a focus of recent chemotaxis research (9–11). Here we demonstrate how the cell’s ability to sense a nutrient depends on the expression levels of the proteins involved in the uptake of that nutrient. Therefore, a molecular-level understanding of the interplay between sensing and uptake is needed to predict chemotactic responses.

A mechanistic understanding of bacterial chemotaxis has developed over decades of research on model organisms, particularly *Escherichia coli*. An *E. coli* cell swims with a “run-and-tumble” pattern, swimming straight before randomly re-orienting by transiently switching the direction of rotation of the flagellar motors (2). The durations of the runs are controlled by a signal-transduction pathway so that attractant binding events at the membrane-bound receptors inhibit motor switches. An additional pathway links rates of methylation and demethylation of the chemoreceptors to the rate of attractant binding events, allowing the cell to store a short-term memory of past measurements and to adapt to background levels of attractant (12–17). The cell thus senses a gradient by measuring the attractant concentration over time and performs chemotaxis by modifying the probability of the next tumbling event in response to the gradient. This behavior has been accounted for in numerous models of chemotaxis. For example, the agent-based Signaling Pathway-based *E. coli* Chemotaxis Simulator (SPECS) incorporates a molecular-level model of chemotaxis to predict the population-level response of cells in environments with gradients of aspartate (18).

Yet existing models do not capture some fundamental cases of chemotaxis, for example when the cell’s uptake of the attractant limits that attractant’s periplasmic concentration, as is the case for *E. coli*’s uptake of the sugar maltose. *E. coli’*s chemotactic response to maltose presents a puzzle because, while all other known *E. coli* sugar chemoattractants are sensed by the minor receptor Trg (19), maltose is sensed by the more abundant aspartate receptor, Tar. Maltose and aspartate are sensed independently by Tar because they can bind simultaneously to distinct sites on the receptor (20), so Tar effectively acts as two distinct receptors sharing the same methylation state. Yet, while *E. coli* can respond to concentrations of aspartate over a few orders of magnitude (21), the dynamic range of *E. coli’s* response to maltose spans just one order of magnitude (19, 22).

Previous work argued that the narrow dynamic range of maltose sensing in *E. coli* mirrors its narrow dynamic range of sensing of other sugars because the sugars bind indirectly to their cognate receptor (19). While aspartate binds directly to Tar, maltose binds to Tar only when in complex with the maltose binding protein, MalE (22, 23). Neumann and coworkers proposed an indirect-binding chemotaxis model to account for the effects of the binding protein (19). However, their model assumes that the concentration of free sugars in the cell periplasm is equal to that in the external environment, which is not the case for maltose chemotaxis (24).

The discrepancy between periplasmic and extracellular concentrations of maltose is due to the relatively large size of the maltose molecule. Under approximately steady-state conditions, the periplasmic concentration of a free substrate is equal to its external concentration when the maximal rate at which the substrate can diffuse into the periplasm (that is, the rate of diffusion when no substrate is present in the periplasm) is greater than the rate of uptake of the substrate into the cytoplasm. This is not necessarily the case for saccharides, which diffuse through general porins at rates that are orders of magnitude lower than those of smaller molecules, such as amino acids (25). Because slow rates of diffusion limit uptake (26), bacteria have evolved specialized porins that facilitate the transport of specific sugars into the periplasm (27). Thus, a cell can regulate the abundance of a particular sugar in its periplasm by regulating the expression of the corresponding porin. This in turn allows the cell to regulate its chemotactic sensitivity to a sugar by altering the amount of that sugar available for a receptor to sense (28, 29).

Previous experiments showed that at extracellular concentrations of approximately 1 μM of maltose, the total rate of maltose uptake into the cytoplasm decreased proportionally with decreasing abundance of the specific maltose porin, LamB (24, 30). Thus, in the micromolar regime, transport is porin-limited. In addition, because of very high concentrations of the maltose binding protein MalE, which scavenges for maltose in the periplasm, the great majority of maltose in the periplasm is bound rather than free. Therefore, when maltose is present in micromolar concentrations in the environment, there are estimated to be significantly lower concentrations of free maltose within the periplasm (24, 26, 30–34). Indeed, Tan and coworkers recently suggested that the periplasmic concentration of free maltose must be lower than the extracellular concentration, based on a discrepancy they found when fitting data of *E. coli* capillary assays to their population-level chemotaxis model, which accounted for indirect binding (35).

To explore the implications of porin-limited transport on chemotaxis, here we develop a detailed molecular-level chemotaxis model that fully accounts for transport dynamics. The model explicitly accounts for the transport of maltose into the periplasm via the maltoporin LamB (28, 36), and out of the periplasm and into the cytoplasm via the ABC maltose transporter MalFGK_2_ (33) (Figure 1). Our transport-and-sensing chemotaxis model incorporates the impact of variable porin and transporter expression on the chemotactic response. Importantly, due to the limiting concentrations of free maltose in the periplasm, our model removes the common simplifying assumption that the free periplasmic concentration of the sensed substrate is independent of the abundance of chemoreceptors (19, 35, 37) and instead explicitly considers how the chemotactic signal is a function of chemoreceptor abundance.

**Figure 1:**
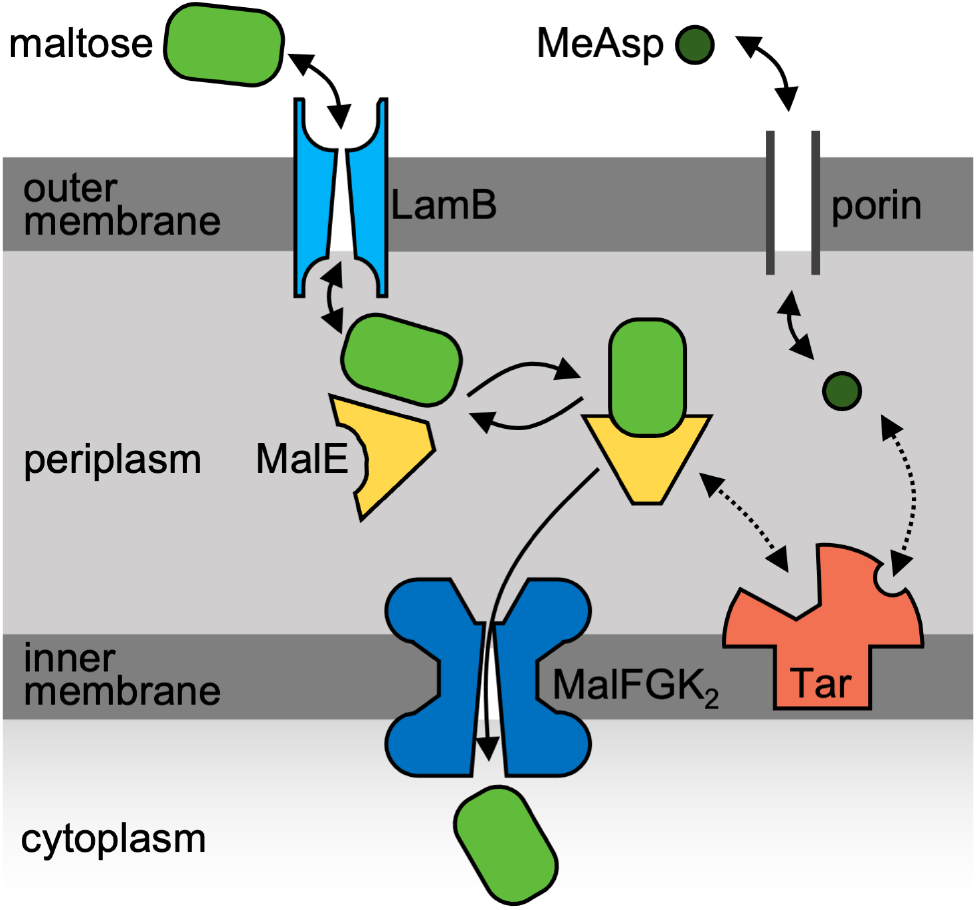
Schematic of the transport and sensing of maltose and MeAsp. MeAsp (α-methyl-DL-aspartate, an analog of the amino acid aspartate) enters and exits the periplasm via general diffusive porins, binds directly to the aspartate receptor, Tar, and is not transported into the cytoplasm. Maltose, in contrast, is larger and enters the periplasm via facilitated diffusion by the maltoporin, LamB. The maltose-binding protein (MalE) both (*i*) binds to the maltose ABC transporter, MalFGK_2_, to allow maltose to be transported into the cytoplasm and (*ii*) allows Tar to sense maltose via a binding site independent of that for MeAsp.

To benchmark our model, we fit it separately to both previous chemotaxis FRET assays (19) and population-level microfluidic experiments that we conducted to quantify *E. coli*’s behavior in gradients of the sugar maltose, either by itself or in opposing gradients of α-methyl-DL-aspartate (MeAsp), a non-metabolizable analog of aspartate. Our model indicates that *E. coli*’s narrow dynamic range of maltose sensing is the result of a trade-off between sensing and uptake. The use of binding proteins allows the cell to achieve high-affinity transport (34, 38) but severely limits the concentration of free maltose in the periplasm so that, to sense maltose gradients, the receptor must bind to the maltose-binding protein complex rather than to maltose alone. Hence, the use of binding proteins tightly couples sensing with uptake so that, if transport were not porin-limited, the chemotactic sensing range would be dictated by the high affinity of the ABC transport system. By instead limiting porin abundance, *E. coli* makes the chemotactic response less sensitive to variations in binding-protein abundance and decouples the regimes at which sensing and cytoplasmic transport saturate. Therefore, although porin-limited transport decreases the total uptake rate at low maltose concentrations, it prevents high-affinity saturation of the chemotactic signal, enabling the cell to sensitively sense micromolar gradients of maltose.

## Results

### A transport-and-sensing chemotaxis model

Because previous research demonstrated that maltoporin abundance can limit the periplasmic concentration of free maltose, we developed a transport-and-sensing model of chemotaxis that accounts for the transport kinetics of maltose into and out of the cell’s periplasm and for the impact of these kinetics on *E. coli’s* sensing of maltose (Figure 1). We provide an outline of the model here, leaving a detailed derivation for Supplemental Appendices 1&2.

During chemotaxis, the instantaneous probability that a cell tumbles is a function of the receptor activity level. We follow previous models and assume that the receptors within a cluster are either all active or all inactive and that the average activity of the clusters, ⟨*a*⟩, is the probability that a receptor cluster is active (37). This probability is a function of the free-energy difference *f* between the active and inactive states. Following the Heterogeneous Monod-Wyman-Changeux (HMWC) model, which describes the integration of multiple chemotactic signals that share the same methylation dynamics (39–42), we assume that the free-energy differences of the receptor types are additive so that

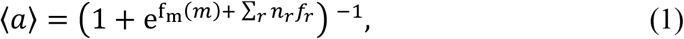

where *f*_*m*_(*m*) is the free-energy difference between an active and inactive receptor cluster in the absence of chemoattractants and depends on the average methylation level of the cluster, *m*; *n*_*r*_ is the number of type-*r* receptors in a cluster; and *f*_*r*_ is the free-energy difference between an active and inactive bound receptor of type *r* (Tu, 2013).

The free-energy difference has the following general form (Tu, 2013):

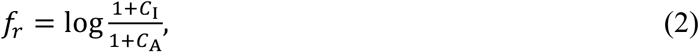

where C_I_ (C_A_) is the ratio of the probabilities of an inactive (respectively, active) receptor being bound versus unbound. For the case of indirect binding, these ratios are

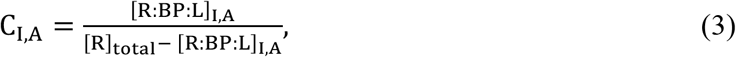

where [R]_total_ is the total effective concentration of the tightly clustered receptors in the periplasm; and [R: BP: L]_I,A_ is the total concentration of ligand–binding protein complex bound to an inactive (or active) receptor.

The quantity [R: BP: L]_I,A_ depends on the concentration of bound maltose-MalE complex available for the receptors to bind, [L: BP]. We assume that this concentration is purely a function of the total concentration of binding proteins in the periplasm, [BP]_total_; the periplasmic concentration of maltose, [L]_p_; and the dissociation constant of the two compounds, *K*_*BP*_:

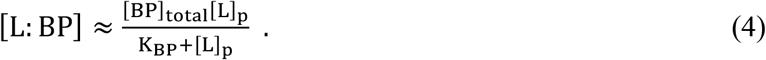

We solve for [L]_p_ as a function of [L]_ext_ by assuming a quasi-steady-state equilibrium in which the rate of maltose transport into the periplasm, *v*_p_, is equal to the rate of its transport out of the periplasm and into the cytoplasm, *v*_c_, where

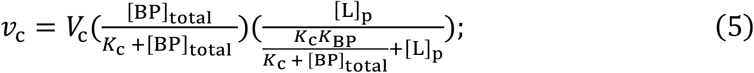

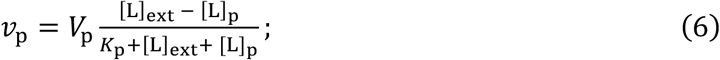

*V*_*c*_ (*V*_p_) is the maximal cytoplasmic (periplasmic) uptake rate, which is a function of the number of expressed ABC transporters (porins); *K*_p_ is the half-saturation constant of the porin; and *K*_c_ is the dissociation constant between the maltose-MalE complex and the ABC transporter.

As a result of our assumption of low periplasmic concentrations of maltose, we cannot make the common simplifying assumption that the free periplasmic chemoattractant concentration is independent of the abundance of chemoreceptors (19, 35, 37). We instead assume that the concentration of bound receptors is

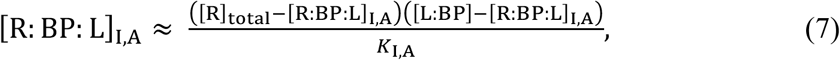

where *K*_I,A_ are the dissociation constants between the inactive (or active) receptor, R, and the maltose-binding protein complex, L: BP. We expand Eq. 7 to the form of a quadratic equation and take its positive root as the solution for [R: BP: L]_I,A_. As we will find below, it is this new, quadratic functional form for the concentration of bound receptors that enables good fits to the data.

The transport-and-sensing model solves for the free-energy difference (Eq. 1) in terms of the substrate transport kinetics and abundances of substrate, receptors, and binding proteins (Eqs. 2–7). In comparison with the indirect-binding chemotaxis model (19), the transport-and-sensing model introduces four new parameters, which describe the Michaelis-Menten kinetics of the transport of maltose into (*K*_p_, *V*_p_) and out of (*K*_c_, *V*_c_) the periplasm. Based on prior observations, we set *K*_p_ = 10 mM (43), *K*_c_ = 100 μM (44), and K_BP_ = 2 μM (32). We take as the free parameters the unknown receptor binding constants, *K*_I_ and *K*_A_, as well as the parameters that may potentially vary with cellular regulation, [R]_total_, [BP]_total_, and *V*_c_/*V*_p_. To obtain a predictive model of maltose chemotaxis, we fit these five free parameters to experimental data.

### E. coli’s population-level chemotactic response to maltose

We conducted chemotaxis experiments using a three-channel microfluidic device. The device creates steady, linearly varying concentrations of one or more chemoattractants within a central test channel by flowing two distinct concentrations of the chemoattractants in two source channels located on either side of the test channel and relying on diffusion through the hydrogel barrier in between (45, 46) (Figure 2, Methods). To create environments with linear concentration profiles of maltose, we placed a buffer solution with no maltose in the right source channel and a solution containing maltose in the left source channel.

**Figure 2:**
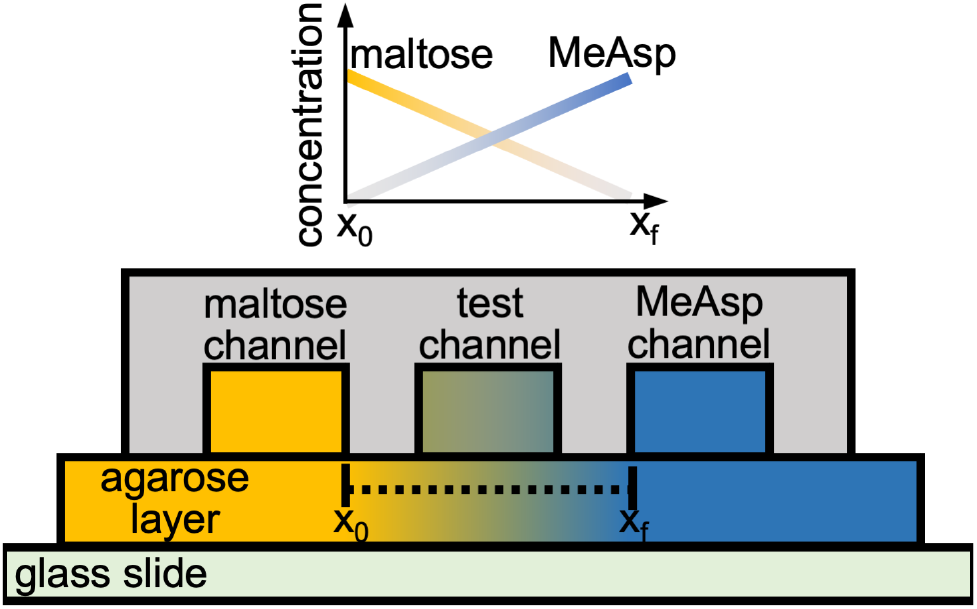
Side view of the three-channel microfluidic device used to expose cells to constant chemoattractant gradients. The channels, fabricated in solute-impermeable polydimethylsiloxane (PDMS), were 20 mm long, 100 μm deep, 600 μm wide, and with 400 μm spacing between each channel. The agarose layer was 0.5 mm thick. Motility medium with a constant maltose concentration was flowed through the left outer channel, and motility medium with a constant MeAsp concentration was flowed through the right outer channel. For the opposing-gradient experiments, both the maltose and MeAsp concentrations were nonzero; for the single-gradient experiments, one of the two outer channels contained only motility medium. The chemoattractants diffused through the agarose layer and up into the test channel, creating two steady linear concentration profiles in opposite directions. Figure adapted from Yawata et al., 2014.

To simplify population-level modeling of the experimental observations, we took several precautions with our experiments. We ensured that the cell densities were sufficiently low and media flow rate sufficiently high so that maltose consumption by the cells was negligible (Methods). We also ensured that the maltose binding protein (MalE) concentrations did not vary over the experiments, as the *mal* regulon is induced by maltose (47). Indeed, a Western blot analysis of MalE showed no difference in the population-averaged MalE expression levels over the range of maltose concentrations tested for the short time durations of our experiments (Figure S1; Methods).

The cells showed a measurable chemotactic response within a narrow range of maltose concentrations created from approximately 0.2 to 20 μM of maltose in the left source channel. The cells showed a strong peak response at 4 μM of maltose (Figure 3*A*; Methods).

**Figure 3:**
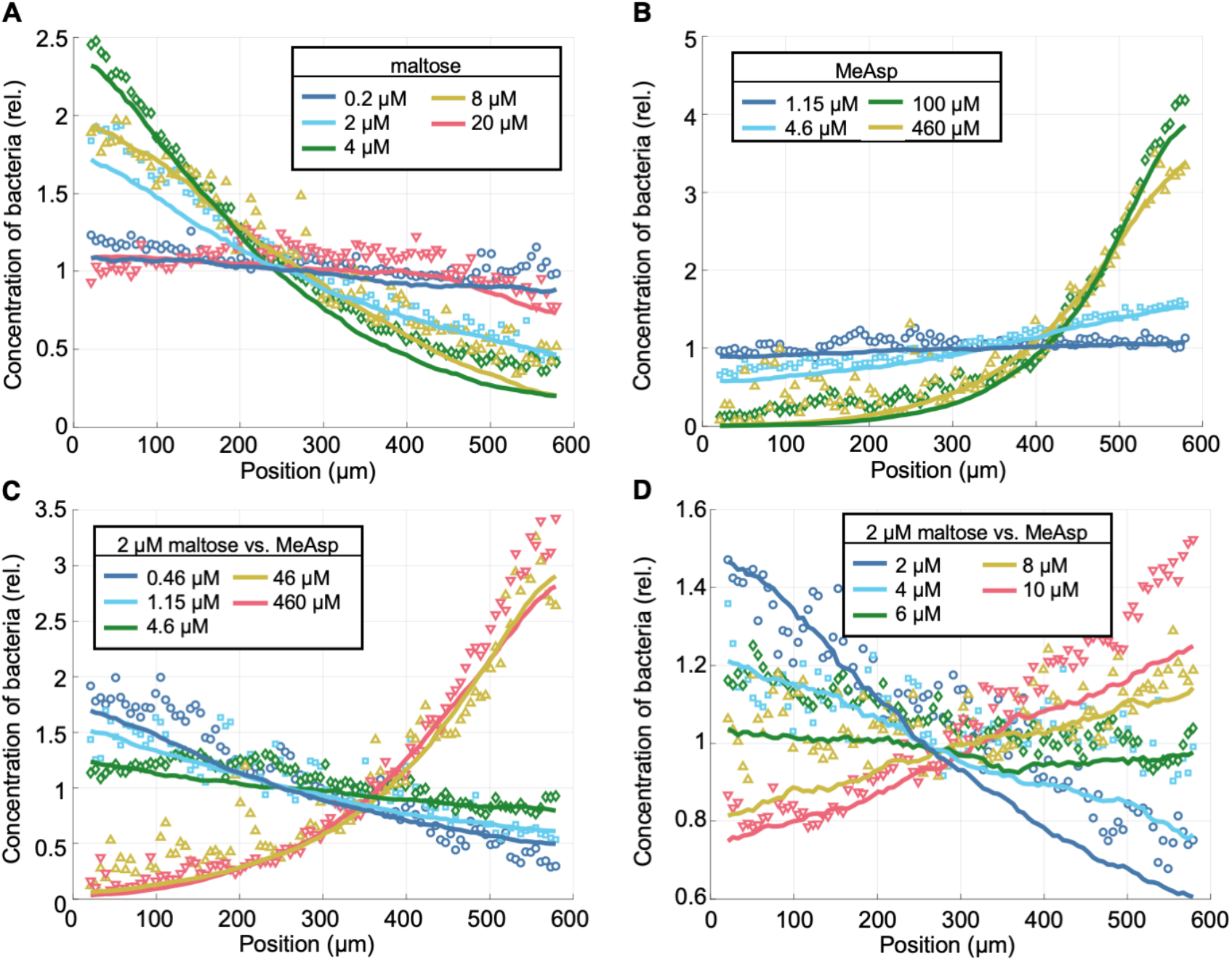
Steady-state distributions from experimental chemotaxis assays in single and opposing gradients of MeAsp and maltose along with the best fits obtained from parameter sweeps of SPECS with the transport-and-sensing model. Each experimental distribution of the relative concentration of bacteria was obtained using the bacterial positions measured over all channel locations and biological replicates (Methods). In each plot, the maximal maltose concentration is at position x = 0 μm and the maximal MeAsp concentration is at position x = 600 μm. Legends give the chemoattractant concentrations in the outer channels. (**A**) Single-gradient maltose experiments. (**B**) Single-gradient MeAsp experiments. (**C**) Opposing-gradient experiments with 2 μM of maltose in the left outer channel and various MeAsp concentrations in the right outer channel. (**D**) Additional opposing-gradient experiments with 2 μM of maltose in the left outer channel and further intermediate MeAsp concentrations in the right outer channel.

To help constrain our model fits, we further measured the chemotactic response in gradients of MeAsp and opposing gradients of maltose and MeAsp, by replacing the buffer solution in the right source channel with a solution containing MeAsp. The response in single gradients of MeAsp match previous assays (Figure 3*B)* (48). In the opposing gradients, depending on the concentrations of the two chemoattractants in these opposing-gradient environments, the cells either favored one substrate over the other and accumulated on one side of the test channel, or were indifferent to the combination of opposing gradients and had a flat distribution across the test channel (Figures 3*C,D*). Under the culture conditions tested, a flat distribution was observed in opposing gradients created by 2 μM of maltose and approximately 6-8 μM of MeAsp.

### Fitting the model to molecular-level data

We first fit the transport-and-sensing model directly to molecular-level data from previous FRET assays of the *E. coli* LJ110 strain (19). For this fitting, we used the data from the attractant dose response as well as of the dynamic range in response to three-fold steps of attractant additions (Methods, Figure S10). We found two different parameter fits that can well describe the FRET assays. One fit predicts protein abundances that match estimates obtained from various previous experimental measurements (Figure S10). However, similar to the original indirect-binding model fitted to the FRET data (19), this fit predicts that the dissociation constants of the bound maltose binding protein to active and inactive Tar (*K*_I(A),Mal_) are in the millimolar regime, whereas direct measurements of the dissociation constants demonstrated that their values are in the micromolar regime (49).

The second obtained fit does achieve dissociation constants that more closely match the measured micromolar values (49) but predicts binding protein abundances of approximately 100 μM, a factor of ten lower than previous estimates (23, 47). It is possible that the maltose binding-protein concentrations were indeed on the order of 100 μM for the strain and culture conditions used in the FRET assays, as this hypothesis is supported by additional FRET assays that showed an increase in chemotactic sensitivity with an increase in binding-protein abundance from an inducible plasmid (19). Our model predicts that increasing binding-protein concentration from a baseline value of 100 μM does indeed increase chemotactic sensitivity (Figure S11). We therefore conclude that our transport-and-sensing model is consistent with the molecular-level FRET data.

However, we were unable to use these molecular-level fits to directly predict the population-level response of our chemotaxis assays because of a crucial discrepancy: the FRET assays show a larger dynamic sensing range than our chemotaxis assays (Figures 3 & S12). We hypothesize that this may be due to differences in experimental conditions or to differences between strain LJ110 used for the FRET assays and strain RP437 used for our chemotaxis assays. Therefore, we fit our transport-and-sensing model directly to the chemotaxis assays.

### Predicting population-level response from a molecular-level understanding

We attempted to fit both the previous indirect-binding model and our transport-and-sensing model directly to the observed population-level responses by incorporating these molecular-level models into a modified version of the agent-based simulator SPECS (18). This modified version accounts for multiple chemoattractant gradients (40, 42) and allows the methylation level to saturate to capture imperfect adaptation (19) (Supplemental Appendix 2, Methods). Due to the intractability of running agent-based simulations within an optimization program, we performed series of parameter sweeps to find good fits (Methods). For the transport-and-sensing model, we constrained the parameter ranges based on estimates from previous literature, whereas for the indirect-binding model we conducted our search over various series of large sweeps. When fitting the indirect-binding model to our single-gradient maltose chemotaxis assays, we found no choice of parameter values that could capture both the narrow range of maltose sensing and the strong peak response at 4 μM of maltose (Figure S2; Methods).

On the other hand, we were able to find a good fit to our chemotaxis assays using our transport-and-sensing model (Figure 3; Methods). We found that our transport-and-sensing model captures not only the steady-state cell distributions in the maltose single-gradient chemotaxis assays but also the steady-state cell distributions in the MeAsp experiments and in the opposing-gradient experiments, both of which we conducted to help us constrain our maltose parameter fits. This supports the validity of our model’s assumption that transport is porin-limited in the chemotactic regime.

Because our model is underdetermined, we obtained our fit by constraining the receptor cooperativity to n_Tar_ = 4, matching the finding that n_Tar_ = 6 for *E. coli* strain LJ110 under similar culture conditions and that the Tar to serine receptor ratio is 1.5 times higher in strain LJ110 than in the strain we used, RP437 (19). In addition, to make the parameter sweep tractable, we used literature values for [R]_total_ = 20 μM (50) and [BP]_total_ = 1 mM (23). With the three parameters thus constrained, our obtained fit is: *K*_I,MeAsp_ = 26 μM, *K*_A,MeAsp_ = 260 μM, *K*_I,Mal_ = 12 μM, *K*_A,Mal_ = 18 μM, and V_c_/V_p_ = 7.5 × 10^−4^ (Figure 3). This fit supports our constraint that n_Tar_ = 4, as the dissociation constants of MeAsp to Tar closely match previous estimates of *K*_I,MeAsp_ = 30 μM and K_A,MeAsp_ = 500 μM (19) as well as our own fit from the FRET data of *K*_I,MeAsp_ = 27.5 μM and K_A,MeAsp_ = 365 μM (Figure S10). Additionally, our estimates for the receptor binding constants (*K*_I,Mal_ = 12 μM and *K*_A,Mal_ = 18 μM) reasonably match observations that these dissociation constants are in the micromolar range (49). Finally, our fit’s estimate that *V*_c_/*V*_p_ = 7.5× 10^−4^ is in line with a rough estimate we obtained previously by fitting our maltose transport model to past experiments that *V*_c_/*V*_p_ ≈ 1× 10^−4^ (34). The consistency of all fitted parameter values with previous observations supports the validity of our transport-and-sensing model. For the obtained fit and taking [L]_ext_ = 1 μM, our model predicts that [L: BP] = 14.9 μM and [L]_p_ = 30.1 nM (Figures 4*A,B*). This predicted nanomolar concentration of free maltose in the periplasm furthermore supports our understanding that maltose transport into the cell is severely porin-limited in the chemotactic regime.

**Figure 4:**
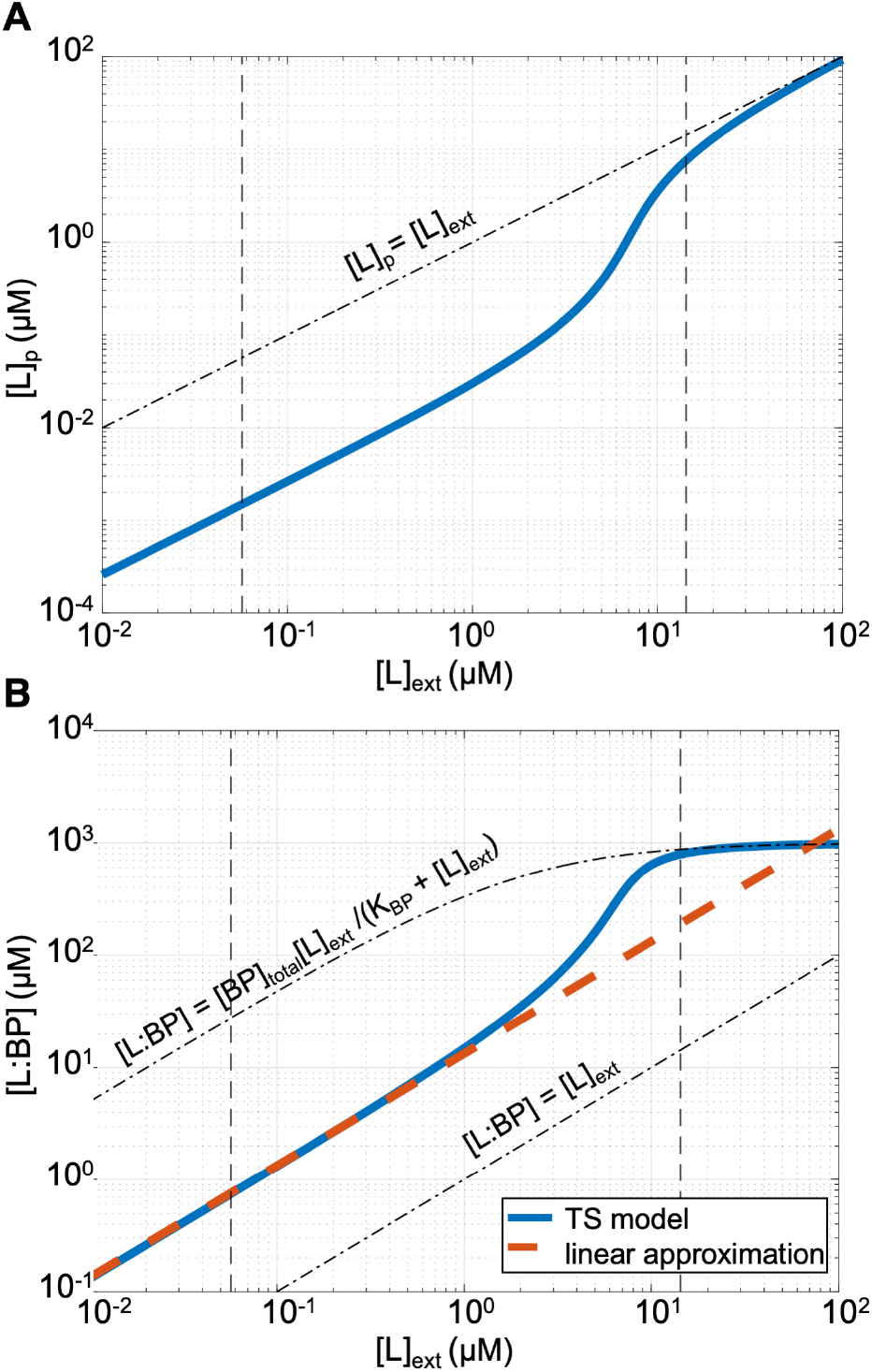
Maltose transport. The transport-and-sensing (TS) model’s predictions of (**A**) periplasmic free maltose concentration, [L]_p_, and (**B**) maltose-binding protein complex concentration, [L:BP], as a function of extracellular maltose concentration using best fit of chemotaxis assays from parameter sweep. Vertical dashed lines show the minimum and maximum maltose concentrations experienced by cells in our experiments. Although the chemotactic signal [L:BP] has a sigmoidal form, it is linear for the majority of the chemotactic sensing range. Thus, we obtain good fits for the observed chemotactic response using the linear approximation (red dashed line) [L:BP] ≈ *α*[L]_ext_ with *α* =: (*K*_c_/*K*_p_)(*V*_p_/*V*_c_) = 13.3. Note that [L:BP] is about an order of magnitude higher than the external maltose concentration [L]_ext_ in the chemotactic regime and approaches [L: BP] ≈ [BP]_total_[L]_ext_/(*K*_BP_ + [L]_ext_) at high external concentrations, where transport is no longer porin-limited so [L]_p_ = [L]_ext_.

Because the transport-and-sensing model has one more free parameter than the indirect-binding model (five instead of four), it may not be surprising that it attains better fits (Figures 3 & S2). However, we find that a modified version of the transport-and-sensing model with only three fitting parameters also captures most of the observed chemotactic response. Because our model predicts a linear relationship between the extracellular maltose concentration [L]_ext_ and the signal [L: BP] for the majority of the maltose sensing range (Figure 4*B*), we additionally fit our data assuming [L: BP] = *α*[L]_ext_ so that there are only three free parameters: *K*_I_/α, *K*_A_/α, and [R]_total_/α (Supplemental Appendix 1). The reduced model captures the observed responses, except for the saturation of the response at 20 μM maltose that is due to the, in fact, sigmoidal relationship between [L: BP] and [L]_ext_ (Figures 4B & S3).

The ability of our reduced model to capture most of the chemotactic response shows that the improved fit of our full model is not due to increased degrees of freedom. Instead, the predictive power of the transport-and-sensing model is due to the new form of the model, specifically the quadratic form of the equation used to solve for the concentration of bound receptors (Eq. 7). This new form accounts for the dependence of limiting free attractant concentrations on receptor abundance and thereby demonstrates the importance of accounting for the relative rates of periplasmic and cytoplasmic transport to predict the chemotactic response.

### A tug-of-war between sensing and uptake explains E. coli’s narrow sensing range for maltose

Our fitted maltose transport model (Eqs. 4-6) indicates that maltose uptake becomes severely porin-limited (i.e., [L]_p_ ≪ [L]_ext_) for extracellular maltose concentrations [L]_ext_⪅ 5 μM (Figure 4*A*), corroborating previous experimental work (32, 33). If maltose transport was instead never limited by maltoporin abundance (so that [L]_p_ = [L]_ext_), we predict that, due to the high abundance of binding proteins, the maltose uptake affinity could be approximately one order of magnitude higher (34), thus permitting higher uptake rates at low extracellular maltose concentrations.

However, our chemotaxis model indicates that, if the cell were to thus increase porin abundance to increase affinity, it would lose its ability to sense maltose gradients in the micromolar regime. Keeping all else equal but assuming that transport is not porin-limited (so that [L: BP] ≈ [BP]_total_[L]_ext_/(*K*_BP_ + [L]_ext_); Supplemental Appendix 1), our model indicates that the chemoreceptors would saturate at lower extracellular maltose concentrations so that the dynamic sensing range would shift down and the peak response would occur at ~ 200 nM of maltose rather than at ~4 μM (Figure S4). This occurs because the maltose binding protein MalE is required for both uptake and sensing; thus, the uptake affinity and chemotactic sensitivity are tightly coupled. Therefore, if uptake saturates at lower extracellular concentrations, the chemotactic response saturates at lower concentrations as well.

There is thus a sensing-uptake trade-off. Although higher affinity allows higher uptake rates at low saccharide concentrations, it precludes the ability of bacteria to sense gradients of these nutrients at ecologically relevant micromolar concentrations. This tug-of-war between increasing affinity—by increasing outer-membrane permeability via increased maltoporin expression—and increasing sensing range by decreasing permeability (Figure S5) provides an explanation for *E. coli*’s narrow sensing range for maltose.

### E. coli’s chemotactic response is insensitive to variations in binding protein abundance

To test the ability of our transport-and-sensing model to predict how variable protein expression levels affect the chemotactic response of cells, we performed additional microfluidic experiments to compare the chemotactic response of cells from the original culture conditions—in which cells had no prior exposure to sugars—with the response of cells grown in medium supplemented with 500 μM of maltose. We conducted an opposing-gradient chemotaxis assay using concentrations of maltose (2 μM) and MeAsp (10 μM) for which neither chemoattractant response was saturated (Figures 3*A,B*). Counterintuitively, cells grown in maltose had a lower relative affinity to maltose than cells grown without maltose (Figure 5A). This was especially surprising because cells grown in maltose had greater maltose-binding protein abundances. As quantified by Western blots, the abundances were nearly double the abundances measured when the cells were grown in the original media without maltose (Figure S1). Interestingly, when we grew cells without maltose but harvested in late exponential phase, we found a very similar decrease in affinity for maltose relative to MeAsp (Figure 5A). At higher optical densities Tar levels are elevated (51, 52), and we also measured a similar increase in maltose binding protein for cells grown on maltose and cells grown without maltose but harvested in stationary-phase (Figure S1).

**Figure 5:**
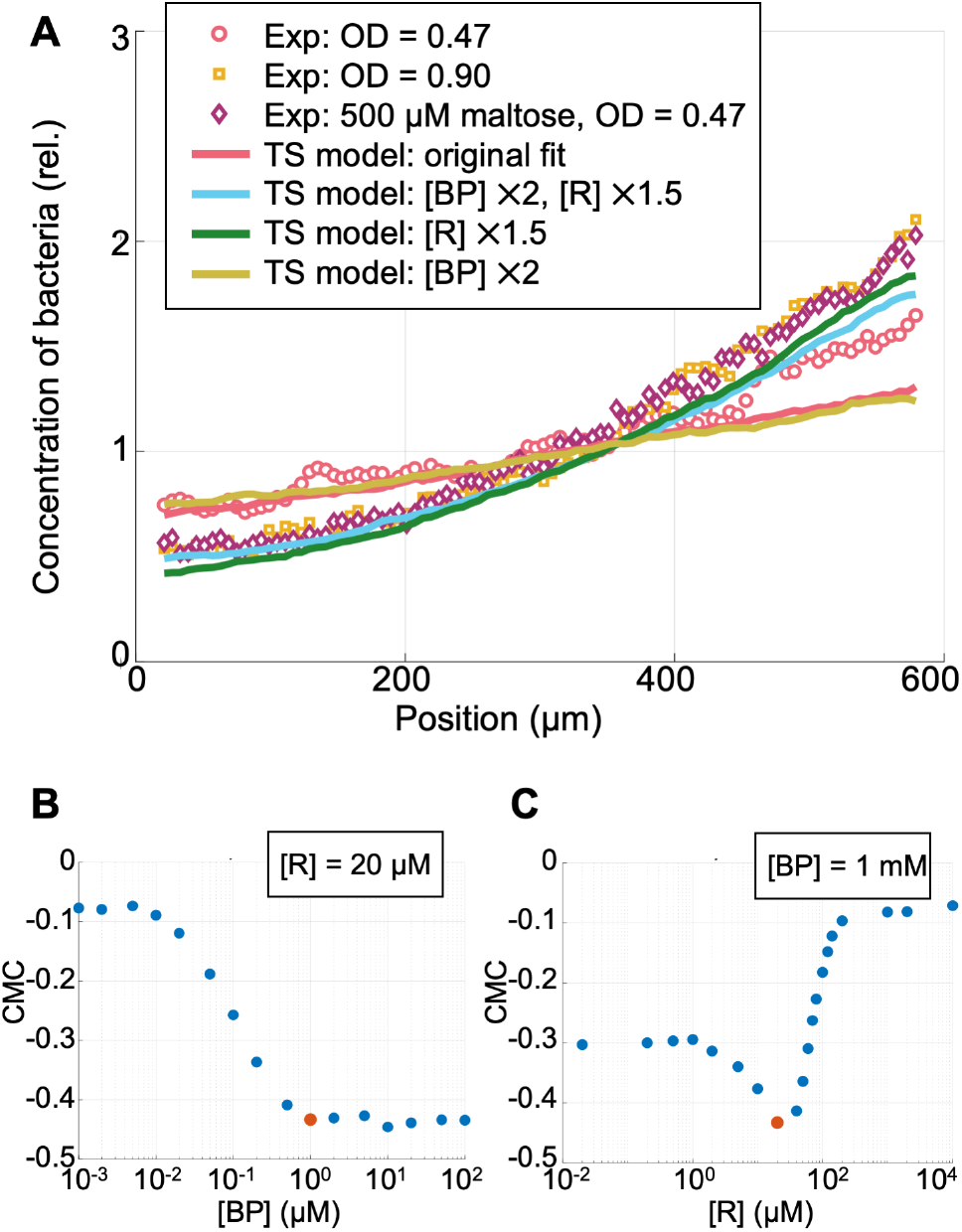
Sensitivity of chemotactic response to variations in binding protein and receptor abundances. (**A**) Steady-state distributions from experimental chemotaxis assays in opposing gradients created using 2 μM maltose and 10 μM MeAsp for cells obtained from different culture conditions. Cells were cultured in tryptone broth and harvested as in other experiments at mid-exponential phase (OD = 0.47), harvested at late-exponential phase (OD = 0.90) when the abundance of Tar is known to be higher, or cultured in tryptone broth supplemented with 500 μM maltose and harvested at OD = 0.47, and then placed in the microfluidic chamber with 2 μM maltose in the left channel and 10 μM MeAsp in the right channel. Curves show fits obtained using an analytical approximation of the transport-and-sensing (TS) model, with variants modifying, by the specified factor, the concentration of MalE, [BP], and the concentration of Tar, [R]. Note that, for increases in [R] by 50%, we also increased n_Tar_ by 50% to a value of n_Tar_ = 6. These results demonstrate that the relative chemotactic abundance is sensitive to Tar abundance but not to binding protein abundance. (**B&C**) We use our fitted SPECS model to predict the peak chemotactic response, at 4 μM of maltose, as a function of both binding protein abundance (with receptor abundance held constant at 20 μM, B) and effective chemoreceptor abundance (with binding protein abundance held constant at 1 mM, C). To quantify the response, we use the chemotaxis migration coefficient (CMC), which is defined as 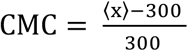, where ⟨x⟩ is the average position in microns of the cells across the 600 μm channel. Our model suggests that the chemotactic response does not vary for sufficiently high binding protein abundances but is very sensitive to variations in receptor abundance. The red dots indicate the responses at 4 μM maltose using our original estimate of the cells’ average binding protein abundance and receptor abundance when grown in tryptone and harvested at OD = 0.47.

The transport-and-sensing model correctly predicts this surprising change in relative affinity, if we allow the parameter values to account for variation in *tar* expression. We found that if we only increase the MalE levels in our model, the distribution did not obtain a good fit (Figure 5A). However, when we assume that supplementing the growth medium with maltose, like harvesting at higher optical densities, increases the expression of Tar by a moderate 50% (51) and also assume that the cell maintains a constant ratio of maximal cytoplasmic and periplasmic uptake rates (*V*_c_/*V*_p_) because *malK* and *lamB* are in the same operon (53), our model provides a good fit of the chemotactic response in opposing gradients (Figure 5A). Thus, although the chemotactic response to MeAsp does not depend on Tar expression levels because the free ligand concentration of MeAsp in the periplasm does not depend on receptor concentration, we found the relative chemotactic affinity to maltose to be highly sensitive to variations in Tar abundance. Yet it was not sensitive to variations in MalE abundance. In fact, our model indicates that the chemotactic response to maltose is independent of binding protein abundance, given that the binding protein abundance is sufficiently high (Figures 5*B*&C, S6&7).

On the other hand, our model predicts that if transport were not porin-limited (so that the periplasmic maltose concentration equals the extracellular concentration), the chemotactic response would be highly sensitive to binding protein abundance (Figures S7 & S8). This stark difference in sensitivity is apparent from the functional forms of the chemotactic signal, [L: BP]. When transport is not porin-limited, [L: BP] is directly proportional to binding protein abundance (Eq. 4; Supplemental Appendix 1). However, in the case of porin limitation (Supplemental Appendix 1, Figure 4*B*), we find

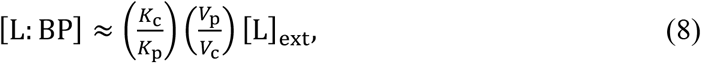

which is independent of binding protein abundance but instead a function of the ratio of abundances of maltoporins, LamB, to maltose ABC transport units, MalFGK_2_.

Because *lamB* and *malK* are adjacent genes in the same operon, we hypothesize that *E. coli* strain RP437 is “hardwired” to maintain a peak chemotactic sensitivity in the micromolar regime. Although protein abundances may vary greatly due to growth conditions and stochasticity, ratios of abundances of proteins expressed from adjacent genes in the same operon vary much less (54). Therefore, in the porin-limited regime, the maltose chemotactic response is robust to variations in maltose transport protein expression levels and is only sensitive to variations in Tar expression levels.

## Discussion

In this work, we developed a transport-and-sensing chemotaxis model, which, unlike previous molecular-level chemotaxis models, does not assume that the periplasmic concentration of a chemoattractant is equal to its extracellular concentration. Rather, it describes the rates of transport of maltose into and out of the periplasm to determine the concentration of maltose-binding protein complex that can bind to receptors. This predictive, mechanistic model captures how *E. coli*’s expression of maltose transport proteins affects its chemotactic response to maltose.

We fit the molecular-level parameters of our model both to previous FRET activity level assays of a mutant *E. coli* strain and also to our population-level, microfluidic chemotaxis experiments. Whereas both the original indirect-binding model and our transport-and-sensing model could explain the FRET data, we found that the indirect-binding model predicted Tar dissociation constants magnitudes higher than previously measured and that only the transport-and-sensing model could fit the FRET data with parameters consistent with previous literature and, in addition, could fit the microfluidic chemotaxis assays in gradients of maltose. This suggests the importance of considering porin-limited transport kinetics to predict population-level chemotactic response.

Previous work argued that, because chemoreception occurs in the periplasm, maltose cytoplasmic transport rates can be drastically reduced without affecting maltose chemotaxis (22, 28). Our work argues the opposite—that the kinetics of cytoplasmic relative to periplasmic transport is crucial to the chemotactic response. This novel understanding allows us to reinterpret previous experimental work, which found that *E. coli* cells with mutations in the MalFGK_2_ transport proteins demonstrated a peak chemotactic response at lower extracellular concentrations of maltose than wild-type cells (22). We conclude that the observed reduction is a direct consequence of the mutants’ reduced cytoplasmic transport rates. The reduced cytoplasmic transport rates increase the concentration of free maltose within the periplasm, causing the binding proteins to saturate at lower extracellular maltose concentrations and thus the chemotactic response to peak at lower maltose concentrations as well.

The transport-and-sensing chemotaxis model provides insight into the roles of the outer membrane and high-affinity transport systems in the chemotactic response of gram-negative bacteria to substrates that do not quickly diffuse through general porins. This is the case for a variety of chemotaxis systems, such as the pathogenic enteric bacteria *Campylobacter jejuni*’s chemotaxis to host glycans (55) and the marine bacteria *Vibrio*’s chemotaxis to chitin (56). Our analysis of the model indicates that, while binding proteins enable high-affinity uptake of a nutrient, they constrain the chemotactic response. Under low nutrient conditions when cells rely on binding proteins for effective transport, the binding proteins capture the majority of the nutrient in the periplasm so that the free concentration is too low to be sensed directly. Therefore, the cell instead senses the nutrient–binding protein complex, and, thus, the uptake affinity is tightly coupled to chemotactic affinity, creating a sensing-uptake trade-off, in which higher uptake affinities result in lower chemotactic sensing ranges.

However, we surprisingly found that, when transport is porin-limited, the chemotactic response is insensitive to small variations in binding protein abundance. Instead, we find that the chemotactic response is highly sensitive to the ratio of porins to transport units. Yet, in the case of *E. coli*’s maltose transport system, these two proteins are transcriptionally co-regulated—which is also a common network motif in the E coli chemotaxis pathway (57)—so their ratio is expected to show little variation. This robustness of the chemotactic response to variations in maltose transport protein expression levels suggests that *E. coli* may have evolved to maintain a maltose sensing range that is independent of transport rates. Thus, we hypothesize that *E. coli* may use maltose not only as a nutrient but also as an environmental cue. It is perhaps for this reason that *E. coli* uses the major receptor Tar rather than the minor receptor Trg to sense maltose separately from all other sugars. However, further work is needed to verify whether *E. coli* cells do, in fact, maintain an invariant sensing range for maltose over a variety of transport conditions.

Our model shows that, while a gram-negative bacterium could use binding protein abundances to tune chemotactic response to growth and environmental conditions if transport were not porin-limited, it can also make chemotactic response independent of transport expression levels by making transport porin-limited. Therefore, although the use of binding proteins limits chemotactic sensitivity and thus creates an uptake-sensing trade-off, it provides a variety of mechanisms for the cell to either independently regulate via protein expression or robustly tune its chemotactic sensing affinity to a particular nutrient based on ecologically relevant conditions.

## Materials and Methods

### Cell culture

We used *E. coli* strain RP437, obtained from the laboratory of JS Parkinson. As controls for the Western blot analyses, we also used two strains from the ASKA library: JW1875-AM (del-tar) and JW3994-AM (del-malE). We grew the cultures overnight in tryptone broth (10 g/L Bacto tryptone, 5 g/L NaCl) in a shaking incubator at 30 °C and 300 rpm, then diluted 1:100 the following morning into fresh tryptone broth. We harvested the cells, unless otherwise noted, when the culture reached OD_600_ = 0.47 in the mid-exponential growth phase. The cells had an exponential growth rate of approximately 0.5 h^−1^. In the one experiment using tryptone broth supplemented with 500 μM of maltose as the growth medium, the cells instead had an exponential growth rate of approximately 0.75 h^−1^. Before their use in experiments, we washed cells twice by centrifuging at 2000 × *g* for 5 min and diluted to OD_600_ = 0.05 in motility medium (10 mM potassium phosphate, 0.1 mM EDTA, 1 μM methionine, 10 mM lactic acid, pH 7) and kept the cells in a 4 °C refrigerator for 30 min.

### Experimental setup

We performed the chemotaxis assays using a microfluidic device made by sandwiching an agarose gel layer between a glass microscope slide and a polydimethylsiloxane (PDMS) layer patterned with three parallel channels. We created the channels by molding the PDMS onto a silicon wafer with positive relief features. We fabricated the device with the following specifications (46): the channels were 20 mm long, 100 μm deep, and 600 μm wide, with 400 μm spacing between each channel, and the agarose layer was 0.5 mm thick and consisted of a 3% (w/v) solution of agarose in motility medium.

Each of the three channels contained an inlet and outlet port. The outer two channels functioned as feeder channels within which a steady flow of media, at a rate of 10 μL per minute, was maintained using a syringe pump (Harvard Apparatus PHD 2000). We set the syringe pump to “refill” mode to create a negative pressure that, along with loosely fitting clips, helped create a seal between the PDMS and agarose. We flowed motility medium with maltose (D-maltose monohydrate; Sigma-Aldrich PHR1497) at concentrations of 0–20 μM in the left channel, and motility medium with MeAsp (α-methyl-DL-aspartate; Sigma-Aldrich M6001) at concentrations of 0–460 μM in the right channel. As molecules diffuse freely through agarose, the flow of these solutions in the outer channels created constant gradients within the agarose and hence within the central test channel (45). For example, with 2 μM of maltose in the left channel and motility medium in the right channel, the cells in the test channel experienced a linear gradient with a slope of approximately 1.4 × 10^−3^ μM/μm and minimum and maximum concentrations of 0.57 μM and 1.43 μM.

Initially, the central channel was empty of liquid, so that, after establishing flow in the outer feeder channels, a steady gradient formed within the lower agarose layer over a timescale of *L*^*2*^/*D*, where *L* is the distance between the two feeder channels and *D* is the diffusivity of the molecules through agarose. The agarose gel layer has a diffusivity very similar to water, so that, for small molecules, *D* ≈ 10^3^ μm^2^/s. Therefore, with a 1,400 μm spacing between the edges of the two outer channels, gradients formed across the agarose layer in about 30 min.

Forty-five minutes after the establishment of flow in the outer channels, we pipetted the refrigerated cells into the test channel and sealed the test channel using glass microscope coverslips. The PDMS and agarose completely blocked advection so that there was no active flow in the central channel; any flow in the central channel was a result of pressure differences between the two ends of the channel and was negligible compared to the swimming speed of the cells. As the channel is only 100 μm deep, the upward diffusion of the chemoattractants from the agarose layer reached a steady state in approximately thirty seconds. Thus, the cells quickly experienced a steady gradient in a no-flow environment. The run-and-tumble chemotaxis of *E. coli* yields an effective diffusivity of *D* ≈ 300 μm^2^/s, so we began data acquisition twenty minutes after the injection of the cells into the test channel, after the bacterial distribution had reached a steady state.

It is important to note that, while MeAsp is non-metabolizable, maltose is metabolizable and is consumed during assays. However, the constant flow of nutrients in the outer channel and the low concentration of cells within the test channel ensured that any changes to maltose concentration within the test channel due to consumption were negligible. Specifically, in the concentration regime of our experiments, *E. coli* cells have an estimated maltose uptake rate of 500 pmoles/min/10^9^ cells (36). There were about 4 ×10^7^ cells/mL in the test channel above an agarose layer 5 times thicker than the channel, which acted as a repository of the nutrient. Therefore, without replacement of maltose, the cells only reduced the concentration within the channel by at most 50 nM after 20 minutes. However, there was in fact replacement, as the wide agarose layer beyond the test channel acted as a source of maltose that was continuously replenished by the flow maintained by the syringe pump.

Under our culture conditions, the cells showed an equal preference for 2 μM of maltose and 6-8 μM of MeAsp in the opposing-gradient experiments (Figure 3*C,D*). For the MeAsp single-gradient experiments, we found the strongest chemotactic response (that is, the steepest steady-state cell distribution in the test channel) when using a MeAsp concentration in the source channel of 46 μM (Figure 3*B*). We were unable to use population-level data to determine the concentration at which Tar’s sensing of MeAsp saturates because the cells are, in fact, repelled by very high concentrations of MeAsp (Figure S9), likely due to pH taxis (58). Therefore, we analyzed the response of cells to MeAsp concentrations up to 500 μM, above which increasing the MeAsp concentration caused a drop in pH (Figure S9*B*).

### Data acquisition and analysis

We acquired images of the chemotaxis assays using a Nikon Eclipse TE2000E inverted microscope fitted with a CCD camera. We imaged the cells using phase contrast with a 20× objective (N.A. = 0.45). For each experiment, we focused the objective mid-depth and took 1-min videos (at 10 frames per second) of at least 5 different 1-mm segments across the entire length of the test channel (obtaining at least 5 technical replicates per biological replicate).

To determine the positions of the bacteria, we analyzed the videos using in-house MATLAB image analysis code that subtracted any non-motile cells. We determined the positions of the bacteria in all frames and thus obtained bacterial position data for 600 frames per segment along the channel. There were about 100 bacteria per frame. We replicated each chemotaxis assay 1–3 times, each time using a new cell culture (obtaining 1-3 biological replicates for each experimental condition tested). We combined the bacterial position data from the technical replicates to obtain a single distribution per biological replicate. Finding very good agreement between the resulting distributions across biological replicates, we summed the bacterial position data over the biological replicates (approximately *N* = 1.2 million bacterial positions) to obtain one distribution corresponding to each experimental condition. This is the distribution we used for all subsequent analysis. Note that, because we caught the same bacterium on multiple frames per channel segment, these *N* positions are not independent.

For the parameter fitting, we first smoothed the data by fitting a power curve to the obtained empirical distributions: for accumulation toward MeAsp, the power fit is of form *f*(*x*) = *ax*^*n*^ + *b*, where *x* is the position in microns along the test channel; for accumulation toward maltose, the power fit is of form *f*(*x*) = *a*(600 − *x*)^*n*^ + *b*. We smoothed the data to subtract noise from poorly swimming and nonmotile cells and chose a power fit because it fitted the distributions well and approximates the analytical expression for the empirical distribution used in our model.

### Agent-based simulations

We ran agent-based simulations of *E. coli* chemotaxis in opposing gradients of maltose and MeAsp. We modified the free-energy difference equations in the original SPECS to incorporate both the HMWC model as well as our transport-and-sensing model. We assumed all additional parameters have the same values as provided in SPECS (18). These parameters include the time discretization, swimming velocity, tumble time, methylation dynamics parameters, Hill coefficient of the motor response, and average directional change due to Brownian rotational diffusion. To describe imperfect adaptation, we modified the methylation dynamics so that the methylation level saturates at a maximum level of 4. In our simulations, when a cell hits the boundary, it modifies its orientation so that it faces away from the boundary with a random angle from a uniform distribution. We used a time step of 0.1 s. To obtain steady-state distributions, we simulated 1,000 cells for eighty minutes of simulated time (48,000 iterations) and averaged their locations over the final forty minutes of the simulated time. (See code on Github.)

### Fitting the model to FRET data

We fit our transport-and-sensing model to previous FRET reporter assays of the dose response and dynamic sensing range of *E. coli* LJ110 to additions of maltose and MeAsp (figures 1A and 2A in Neumann et al., 2010). We used MATLAB’s general nonlinear optimizer, *fmincon*, and solved for the parameter values that minimized the sum of squares of the differences between the observed and predicted dynamic range measurements, constraining the cooperativity of Tar to n_Tar_ = 6. (See our code on Github.)

### Fitting SPECS models to the chemotaxis assays

Due to the computational intractability of running sets of agent-based simulations within an optimization program, we performed parameter sweeps to find good fits for SPECS that used either the original indirect-binding model (19) or our transport-and-sensing model to describe the chemotactic response to maltose.

Because we were unable to find a good fit using the indirect-binding model, we used multiple parameter sweeps—over a large range and with finer discretizations—to ensure that we were not missing a better fit. Over these various sweeps, we spanned: K_BP_ = [0.5: 0.1: 3], p_0_ = [0: 0.05: 0.5], K_I_/[BP] = [0.2: 0.1: 1 2: 2: 50 50: 10: 500], and K_A_/K_I_ = [1.2:0.1:10].

On the other hand, because of the even larger parameter space for our transport-and-sensing model, our approach was to use previous estimates from the literature to constrain the parameter space as much as possible and then show that, even with these constraints, we could find good fits that captured the response of our chemotaxis assays. We constrained the binding protein abundance to [BP]_total_ = 1000 μM (59), the receptor abundance to [R]_total_ = 20 μM (50), and the ratio of the cytoplasmic to periplasmic maximal uptake rates to *V*_c_/*V*_p_ ∈ [10^−5^, 10^−3^] (34). We used estimates that the aspartate to serine receptor ratio (Tar/Tsr) is 1.5 times higher in an LJ110 strain than in an RP437 strain (19) to constrain the receptor cooperativity to n_Tar_ = 4.

To compare fits over our parameter sweeps, we used the following goodness-of-fit measure:

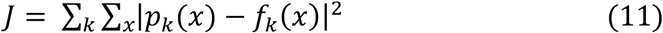

where *f*_*k*_(*x*) (*p*_*k*_(*x*)) is the smoothed empirical distribution from experiment *k* (simulation *k)* and *x* are 100 binned positions spanning the 600 μm channel. To obtain the fits for SPECS with the transport-and-sensing model, we summed over the following single- and opposing-gradient experiments: five maltose single-gradient experiments (0.2, 2, 4, 8, and 20 μM maltose), four MeAsp single-gradient experiments (1.15, 4.6, 46, and 460 μM MeAsp), and eight maltose and MeAsp opposing-gradient experiments (2 μM maltose and 0.46, 2, 4, 6, 8, 10, 46, and 460 μM MeAsp). In out attempt to fit SPECS with the indirect-binding model, we summed only over the five maltose single-gradient experiments.

### Western blot assay of MalE expression

To confirm that there was no change in MalE expression levels over the maltose concentrations and time durations of our experiments, we performed a Western blot analysis of MalE (Figure S1). We cultured, washed, and harvested the cells according to the protocol described above and then placed the cells in 3mL of motility medium with various concentrations of maltose. Because the cells experienced the maltose gradients for at most 30 minutes in our chemotaxis assays, after 30 minutes, we froze the samples at −80 °C until we performed immunoblotting. To lyse the cells, we added 200 μL lysis buffer (50 mM Tris, 100 mM NaCL, 0.1% Triton X-100, 250 U/mL benzonase nuclease, and 0.4 mg/mL lyzosyme) into each 3 mL frozen sample, vortexed, and shook tubes for 30 min at 37 °C. We then added loading Laemmli buffer (1:4) and incubated at 95 °C for 5 min. We loaded 10 μL of each sample into a pre-prepared 12% sodium dodecyl sulfate (SDS) gel (Bio-Rad) and separated the protein by electrophoresis at 100 V for 1 h in a Bio-Rad Tetra cell apparatus. We blotted the gel against a polyvinylidene difluoride (PVDF) membrane using transfer buffer (25 mM Tris, 190 mM glycine, 20% methanol) at 100 V for 1 h. We then blocked the blot using blocking buffer (3% bovine serum albumin in phosphate buffered saline with tween 20 (PBST)) for 1 h at room temperature. For the primary antibody, we used 1 μg/mL anti-MalE (unconjugated rabbit polyclonal antibody; LS Bio LS-C355688) in blocking buffer for 1 h at room temperature, and then placed it in a 4 °C refrigerator overnight. The following morning, we washed the blot three times in PBST. For the secondary antibody, we used 1:5000 goat anti-rabbit IgG secondary antibody (Thermo Fisher 65-6120) in blocking buffer, maintained for 2 h at room temperature and then washed three times in PBST. We stained the blot with 1-step Ultra TMB blotting solution, leaving the blot covered at room temperature. We acquired images after 10, 30, and 90 minutes exposure. We quantified the bands of the resulting blot using ImageJ software (imagej.nih.gov/ij/) and normalized the intensity of each band by their respective total protein concentrations, obtained by absorbance measurements at 280 nm (A280) from Thermo-Scientific NanoDrop UV-Vis spectrophotometry.

## Author Contributions

NN, EF, and RS conceived the idea; NN and FM designed the assay experiments; NN and NL developed the model; YY developed the microfluidic device used for the assays; NN performed the chemotaxis experiments, analyzed the results, and fitted the model to the chemotaxis assays and previous FRET reporter assays; FM, JK, and VF contributed to data analysis; UA designed and performed the Western blot experiments; NN, UA, JK, FM, NL, VF, and RS interpreted the results; and NN, JK, VF, RS, and NL wrote the paper.

## Acknowledgments

The authors acknowledge the support of a Gordon and Betty Moore Foundation Marine Microbiology Initiative Investigator Award (grant 3783 to R.S.) and a grant from the Simons Foundation through the Principles of Microbial Ecosystems (PriME) collaboration (to R.S.). Contributions to the editing of this paper by Dr. Russell Naisbit are gratefully acknowledged. N.N. would also like to thank Professor Ned Wingreen for his invaluable feedback.

## Code availability

The code used for the agent-based simulations and data fitting is available at: https://github.com/noelenorris/maltose_chemotaxis.git

## Notes

https://github.com/noelenorris/maltose_chemotaxis.git

### Competing Interest Statement

The authors have declared no competing interest.

