## Supplemental Material for "Bacterial chemotaxis to saccharides is governed by a trade-off between sensing and uptake"

Noele Norris, Uria Alcolombri, Johannes M. Keegstra, Yutaka Yawata,
Filippo Menolascina, Emilio Frazzoli, Naomi M. Levine, Vicente I. Fernandez, Roman Stocker

### Supplemental Appendix 1: A maltose transport model

To describe the rate of maltose uptake into the cytoplasm, we use an ABC transport model and Michaelis-Menten approximation of ABC transport that we previously derived (Norris et al. 2021). Because it has been shown in *E. coli* that the abundance of the maltose binding protein greatly exceeds the abundance of transporters (Boos and Shuman 1998), the Michaelis-Menten approximation is valid. Therefore, we take the concentration of complex of maltose bound to the maltose binding protein to be:

$$[\mathrm{L{:}BP}] \approx [\mathrm{BP}]_{\mathrm{total}} \frac{[\mathrm{L}]_{\mathrm{p}}}{K_{\mathrm{BP}} + [\mathrm{L}]_{\mathrm{p}}}. \tag{A-1}$$

where $[\mathrm{BP}]_{\mathrm{total}}$ is the total concentration of maltose binding protein in the periplasm; $[\mathrm{L}]_{\mathrm{p}}$ is the concentration of free maltose in the periplasm; and $K_{\mathrm{BP}}$ is the dissociation constant of maltose and maltose binding-protein.

We take the uptake rate of maltose into the cytoplasm to be

$$v_{\mathrm{c}} \approx V_{\mathrm{c}} \frac{[\mathrm{BP}]_{\mathrm{total}}}{K_{\mathrm{c}} + [\mathrm{BP}]_{\mathrm{total}}} \frac{[\mathrm{L}]_{\mathrm{p}}}{\frac{K_{\mathrm{c}} K_{\mathrm{BP}}}{K_{\mathrm{c}} + [\mathrm{BP}]_{\mathrm{total}}} + [\mathrm{L}]_{\mathrm{p}}}, \qquad V_{\mathrm{c}} = k_2 [\mathrm{T}]_{\mathrm{total}}, \tag{A-2}$$

where: $V_{\mathrm{c}}$ is the maximal cytoplasmic uptake rate; $K_{\mathrm{c}}$ is the dissociation constant of the bound maltose-maltose binding protein complex and the transport unit; $k_2$ is the turnover rate of the membrane-bound transport unit; and $[\mathrm{T}]_{\mathrm{total}}$ is the total concentration of transport units in the periplasm.

While transport into the cytoplasm is active and thus can occur against concentration gradients, transport into the periplasm via porins is diffusive. Thus, while the cytoplasmic uptake rate has the above form, the periplasmic uptake rate is better described by the following Michaelis-Menten equation

(Bosdriesz et al. 2018):

$$v_\mathrm{p} \approx V_\mathrm{p}\frac{[\mathrm{L}]_\mathrm{ext} - [\mathrm{L}]_\mathrm{p}}{K_\mathrm{p} + [\mathrm{L}]_\mathrm{ext} + [\mathrm{L}]_\mathrm{p}}, \tag{A-3}$$

where $K_\mathrm{p}$ is the half-saturation constant of the specific porin; and $V_\mathrm{p}$ is the maximal rate of uptake, which is a function of the number of expressed porins.

At steady-state, the periplasmic transport rate ($v_\mathrm{p}$) must be equal to the cytoplasmic transport rate ($v_\mathrm{c}$). Thus, we equate the rates from Equations A-2 and A-3 to solve for $[\mathrm{L}]_\mathrm{p}$ as a function of $[\mathrm{L}]_\mathrm{ext}$.

### A linear approximation

Analysis of the form of [L:BP] as a function of $[\mathrm{L}]_\mathrm{ext}$ for the obtained parameter fits (Figure 4) shows that, for $[\mathrm{L}]_\mathrm{ext} \lessapprox 5\ \mu\mathrm{M}$, [L:BP] can be very well approximated by:

$$[\mathrm{L{:}BP}] = \alpha[\mathrm{L}]_\mathrm{ext}, \qquad \alpha = \frac{K_\mathrm{c}V_\mathrm{p}}{K_\mathrm{p}V_\mathrm{c}}. \tag{A-4}$$

This analytical approximation can be obtained by assuming that $[\mathrm{L}]_\mathrm{p} \ll [\mathrm{L}]_\mathrm{ext}$ and demonstrates that, in the micromolar regime, chemotactic response is independent of binding protein abundance (Figures S7 & S8) and only dependent on the ratio of porin abundance to transport unit abundance.

### Supplemental Appendix 2: A chemotaxis model

To model the chemotactic response of *E. coli* in mixed gradients of maltse and methyl-aspartate, we extend the Signaling Pathway-based *E. coli* Chemotaxis Simulator (SPECS; Jiang et al. 2010) to incorporate: (*i*) the heterogeneous MWC model (Keymer et al. 2006; Mello and Tu 2005) to consider the chemotactic response of cells to multiple chemoattractants; and (*ii*) our new sensing-and-transport model of chemotaxis to maltose that takes into account the transport kinetics of maltose into and out of the periplasm, as well as the indirect binding of maltose to the aspartate receptor via the maltose-binding protein.

Analogous to the derivation of the MWC model presented by Tu (2013), we derive the free energy difference between the active and inactive states of a receptor cluster. We assume that the receptors in a cluster are either all active or all inactive. A single receptor has four possible states: active ($a = 1$) or inactive ($a = 0$) and bound ($l = 1$) or vacant ($l = 0$), with probability, $P(a, l)$, where:

$$\frac{P(1,0)}{P(0,0)} = e^{-f_\mathrm{m}(m)}, \quad \frac{P(0,1)}{P(0,0)} = C_\mathrm{I}, \quad \text{and} \; \frac{P(1,1)}{P(1,0)} = C_\mathrm{A}, \tag{A-5}$$

where $C_\mathrm{I}$ and $C_\mathrm{A}$ are functions that we derive below.

Because the expected activity level of a single receptor is $\langle a \rangle_\text{receptor} = P(1,0) + P(1,1)$ and $P(0,0) + P(0,1) + P(1,0) + P(1,1) = 1$,

$$\langle a \rangle_\text{receptor} = \frac{e^{-f_\mathrm{m}(m)} \left[1 + C_\mathrm{A}\right]}{1 + C_\mathrm{I} + e^{-f_\mathrm{m}(m)} \left(1 + C_\mathrm{A}\right)}. \tag{A-6}$$

We define the free energy difference, $\Delta f$, such that $\langle a \rangle_\text{receptor} = \left(1 + e^{-\Delta f}\right)^{-1}$. Thus,

$$\Delta f = -f_\mathrm{m}(m) - \log\left[\frac{1 + C_\mathrm{I}}{1 + C_\mathrm{A}}\right]. \tag{A-7}$$

Because we assume that all of the $n$ receptors are active or all of them are inactive, the expected activity level of the entire receptor cluster is $\langle a \rangle = \left(1 + e^{-n\Delta f}\right)^{-1}$ (Phillips et al. 2012).

Therefore, a general formulation for the average activity level of a cell sensing chemoattractant L is

$$\langle a \rangle = \frac{1}{1 + e^{[n f_\mathrm{m}(m) + n f_\mathrm{L}([\mathrm{L}])]}}, \quad f_\mathrm{L}([\mathrm{L}]) = \log\left[\frac{1 + C_\mathrm{I}}{1 + C_\mathrm{A}}\right], \tag{A-8}$$

where $m$ is the methylation level and $C_\text{I}$ ($C_\text{A}$) is the ratio of the probabilities of a receptor being bound versus ligand-free for an inactive (respectively, active) receptor.

We first rederive the $C$ term for MeAsp to demonstrate how the MWC model can be extended to describe maltose chemotaxis. By definition, the dissociation constant is $K_\text{MeAsp} = [\text{R}]_\text{free}[\text{MeAsp}]/[\text{R:MeAsp}]$, where $[\text{R}]_\text{free}$ and [MeAsp] are the effective concentrations of free receptor and ligand in the periplasm and [R:MeAsp] is the concentration of bound receptor. Defining $[\text{R}] = [\text{R}]_\text{free} + [\text{R:MeAsp}]$, $[\text{R:MeAsp}] = [\text{R}][\text{MeAsp}]/\left(K_\text{MeAsp} + [\text{MeAsp}]\right)$. Thus, the probability a receptor is bound is

$$P = \frac{[\text{R:MeAsp}]}{[\text{R}]} = \frac{[\text{MeAsp}]}{K_\text{MeAsp} + [\text{MeAsp}]}, \tag{A-9}$$

so

$$C = \frac{P}{1-P} = \frac{[\text{MeAsp}]}{K_\text{MeAsp}}. \tag{A-10}$$

Therefore, distinguishing an active from an inactive receptor,

$$f_\text{MeAsp}([\text{MeAsp}]) = \log\left[\frac{1 + [\text{MeAsp}]/K_\text{I,MeAsp}}{1 + [\text{MeAsp}]/K_\text{A,MeAsp}}\right]. \tag{A-11}$$

Note that because MeAsp is not metabolized by the cell, the steady-state concentration of MeAsp in the periplasm is equal to the extracellular concentration of MeAsp.

Adding sensing to the ABC transport model complicates an already complicated system. To sense maltose, the receptors must compete with the ABC transporters to bind with the ligand-binding protein complex. Optimally, however, sensing would minimally hinder transport to thus minimally decrease the cell's growth rate. We thus make the simplifying assumption that sensing does not affect transport but simply "reads" the state of the system. This is a reasonable approximation given that the abundance of maltose-binding protein greatly exceeds the abundance of the cognate ABC transporter.

Therefore, we assume that the ABC transporters and receptors do not compete for the ligand-binding protein complex and likewise assume that the receptors do not affect binding and dissociation of the binding protein with maltose. Therefore, we modify the transport model to incorporate sensing only via a simple modification to Equation A-1:

$$[\text{L:BP}]_0 \equiv [\text{R:L:BP}] + [\text{L:BP}] \approx [\text{BP}]_\text{total}\frac{[\text{L}]_\text{p}}{K_\text{BP} + [\text{L}]_\text{p}}, \tag{A-12}$$

where R:L:BP is the receptor bound to the ligand-binding protein complex.

We assume that the receptor can only bind to the complex and not to the binding protein on its own so that we can describe sensing by the following mass-action kinetics:

$$\mathrm{R} + \mathrm{L{:}BP} \underset{k_\mathrm{r}}{\overset{k_\mathrm{f}}{\rightleftharpoons}} \mathrm{R{:}L{:}BP}$$

At steady-state, the concentration of the bound receptor does not change, so that

$$[\mathrm{R{:}L{:}BP}] = \frac{[\mathrm{R}][\mathrm{L{:}BP}]}{K}, \quad K = \frac{k_\mathrm{r}}{k_\mathrm{f}}. \tag{A-13}$$

Combining Equations A-12 and A-13, we obtain that the total concentration of maltose-MBP complex bound to an inactive (I) or active (A) receptor is

$$[\mathrm{R{:}L{:}BP}]_\mathrm{I,A} = \frac{\left([\mathrm{R}]_\mathrm{total} - [\mathrm{R{:}L{:}BP}]_\mathrm{I,A}\right)\left([\mathrm{L{:}BP}]_0 - [\mathrm{R{:}L{:}BP}]_\mathrm{I,A}\right)}{K_\mathrm{I,A}}, \tag{A-14}$$

where $K_\mathrm{I,A} = K_\mathrm{I}$ when all of the receptors in the cluster are inactive and $K_\mathrm{I,A} = K_\mathrm{A}$ when all of the receptors are active. Therefore, the terms $[\mathrm{R{:}L{:}BP}]_\mathrm{I,A}$ are the solutions to quadratic equations, and the free energy term for maltose is:

$$f_\mathrm{Mal}([\mathrm{L}]_\mathrm{ext}) = \log\left[\frac{1 + C_\mathrm{I}}{1 + C_\mathrm{A}}\right], \quad C_\mathrm{I,A} = \frac{[\mathrm{R{:}L{:}BP}]_\mathrm{I,A}}{[\mathrm{R}]_\mathrm{total} - [\mathrm{R{:}L{:}BP}]_\mathrm{I,A}}. \tag{A-15}$$

We can use the Heterogeneous MWC (HMWC) model (Keymer et al. 2006; Mello and Tu 2005) to describe the average activity level in mixed gradients of MeAsp and maltose because MeAsp and the maltose-binding protein complex bind independently to distinct sites of Tar (Mowbray and Koshland 1987):

$$\langle a \rangle = \frac{1}{1 + e^{n_\mathrm{Tar}\left[f_\mathrm{m}(m) + f_\mathrm{MeAsp}([\mathrm{MeAsp}]) + f_\mathrm{Mal}([\mathrm{L}]_\mathrm{ext})\right]}}. \tag{A-16}$$

### Supplemental Figures

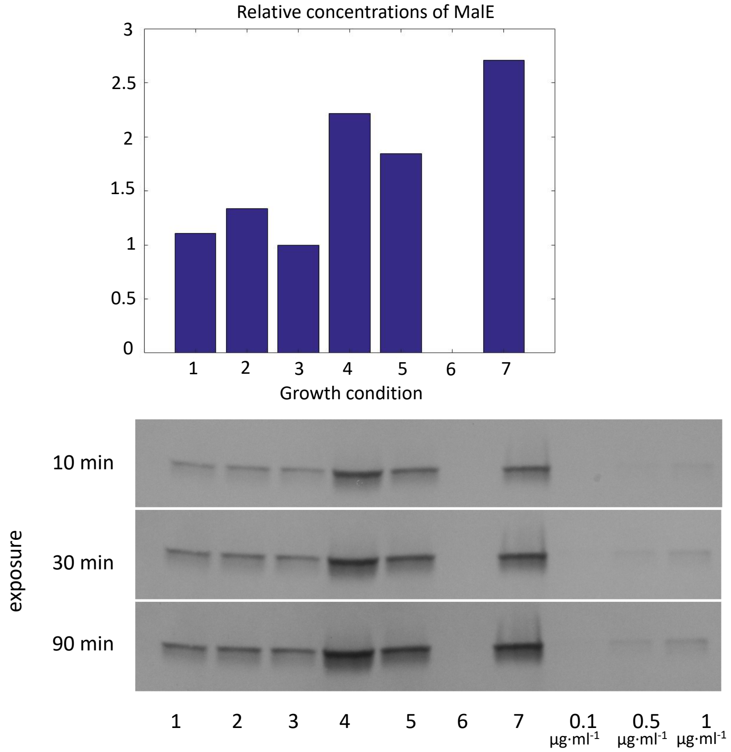


**Figure S1: Western blot for MalE.** The bar plot shows the relative concentration of MalE normalized by total protein concentrations over the following growth and experimental conditions: (1) wild-type cell grown in tryptone broth, put in solution without any maltose, (2) wild-type cell grown in tryptone broth, put in 1 µM maltose, (3) wild-type cell grown in tryptone broth, put in 10 µM maltose, (4) wild-type cell grown in tryptone broth and harvested at $OD_{600}$ = 0.9, put in 1 µM maltose, (5) wild-type cell grown in tryptone broth with 500 µM maltose, put in 1 µM maltose, (6) del-malE strain grown in tryptone broth with 500 µM maltose, put in 1 µM maltose, (7) del-tar strain grown in tryptone broth with 500 µM maltose, put in 1µM maltose. (8-10) three MalE concentrations. The loading volume was 10 µL for all samples. We conclude that MalE abundances were invariant over the experimental conditions shown in Figure 3 because abundances did not vary greatly over lanes 1-3. We estimated that supplementing the tryptone broth with maltose during growth doubled MalE abundance by comparing lanes 1 and 5.

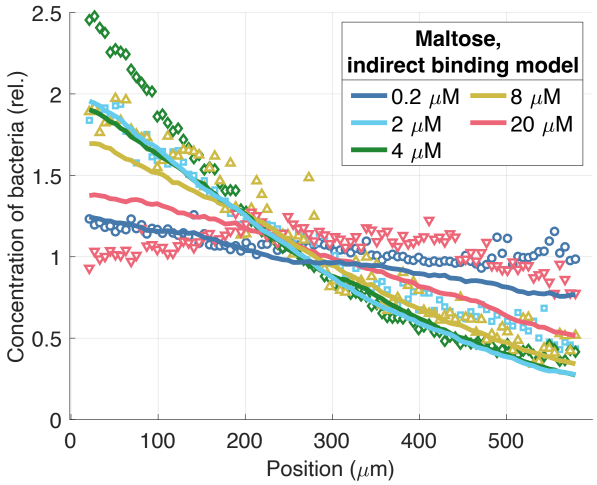


**Figure S2: Predicted steady-state distributions of cells using best fit of SPECS with indirect-binding model.** Experimental cell distributions in maltose with predictions from SPECS simulator incorporated with indirect-binding model and using best-fit parameters from parameter sweep (Methods): $n_{Tar} = 6$, $K_{BP} = 2.6$ μM, $K_{I,Mal}/[BP] = 0.8$, $K_{A,Mal}/[BP] = 1.92$, and $p_0 = 0$.

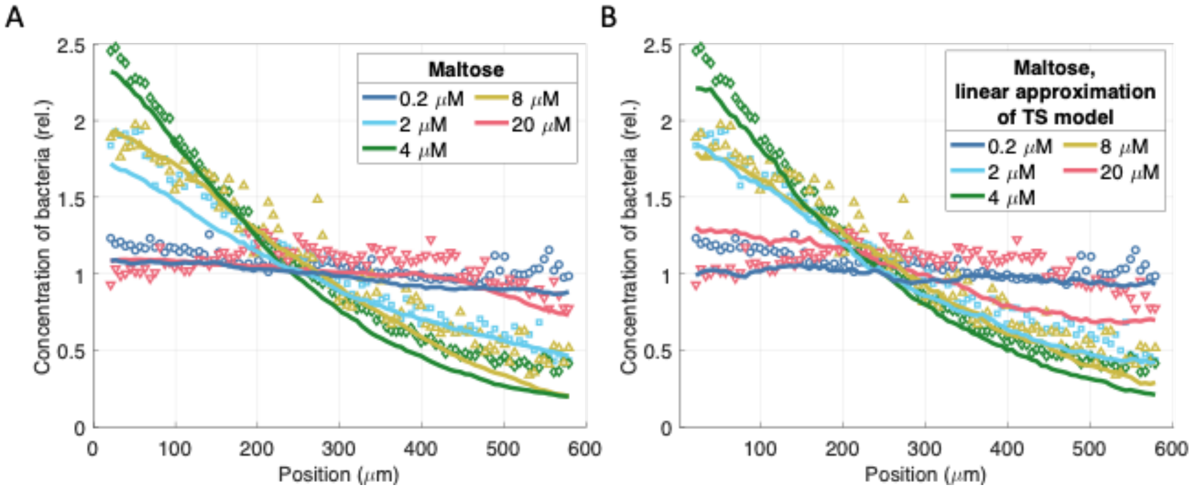


**Figure S3: The linear approximation of the transport-and-sensing model.** (A) Experimental data and best fit using transport-and-sensing chemotaxis model. (B) Experimental data and best fit using linear approximation of transport-and-sensing chemotaxis model, in which we assume $[L{:}BP] \approx \alpha[L]_{ext}$. The best-fit parameter values obtained from the parameter sweep are: $K_I/\alpha =$ 0.72 μM, $K_A/\alpha = 1.18$ μM, and $[R]/\alpha = 1.18$ μM. The linear approximation does not capture the saturation of the response at 20 μM maltose because our transport model suggests that [L:BP] is, in fact, a sigmoidal function of $[L]_{ext}$ (Figure 4).

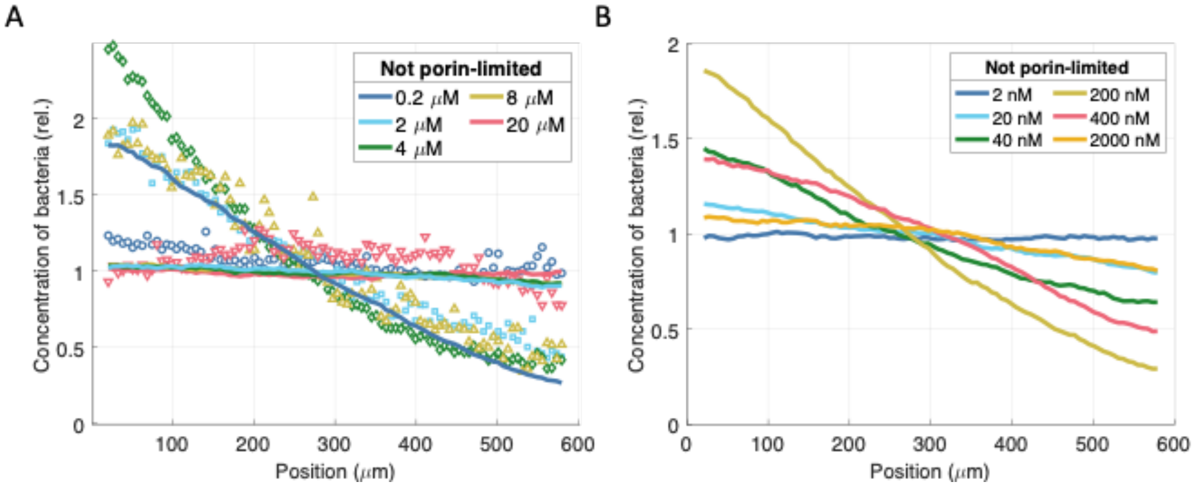


**Figure S4: Chemotactic response if transport were not porin-limited.** We obtained the above plots using the fitted parameters of the transport-and-sensing chemotaxis model but incorrectly assuming that $[L]_p \approx [L]_{ext}$ so that $[L{:}BP] \approx [BP]_{total}[L]_{ext}/(K_{BP} + [L]_{ext})$. (A) We plot the experimental data for reference. When we assume that transport is no longer porin-limited, our model predicts that the cell can no longer sense gradients in the micromolar regime. (B) Instead, its chemotactic sensitivity has shifted down to the nanomolar regime.

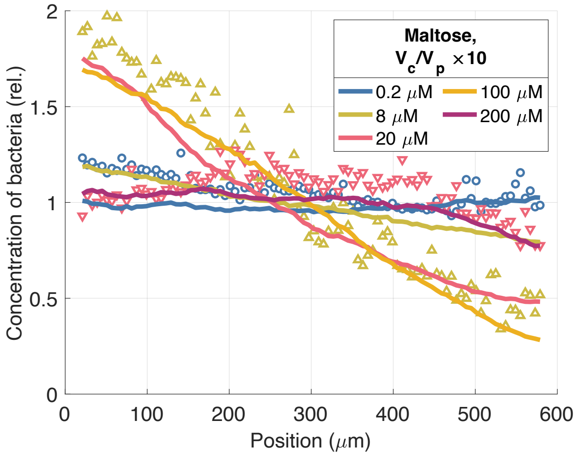


**Figure S5**: **Increasing the dynamic sensing range by decreasing outer-membrane permeability.** If the cell further decreased maltoporin abundance to make transport even more porin-limited, it could increase its dynamic sensing range. However, this would decrease uptake affinity. We hypothesize that this trade-off between sensing range and uptake affinity may explain *E. coli*'s narrow sensing range for maltose.

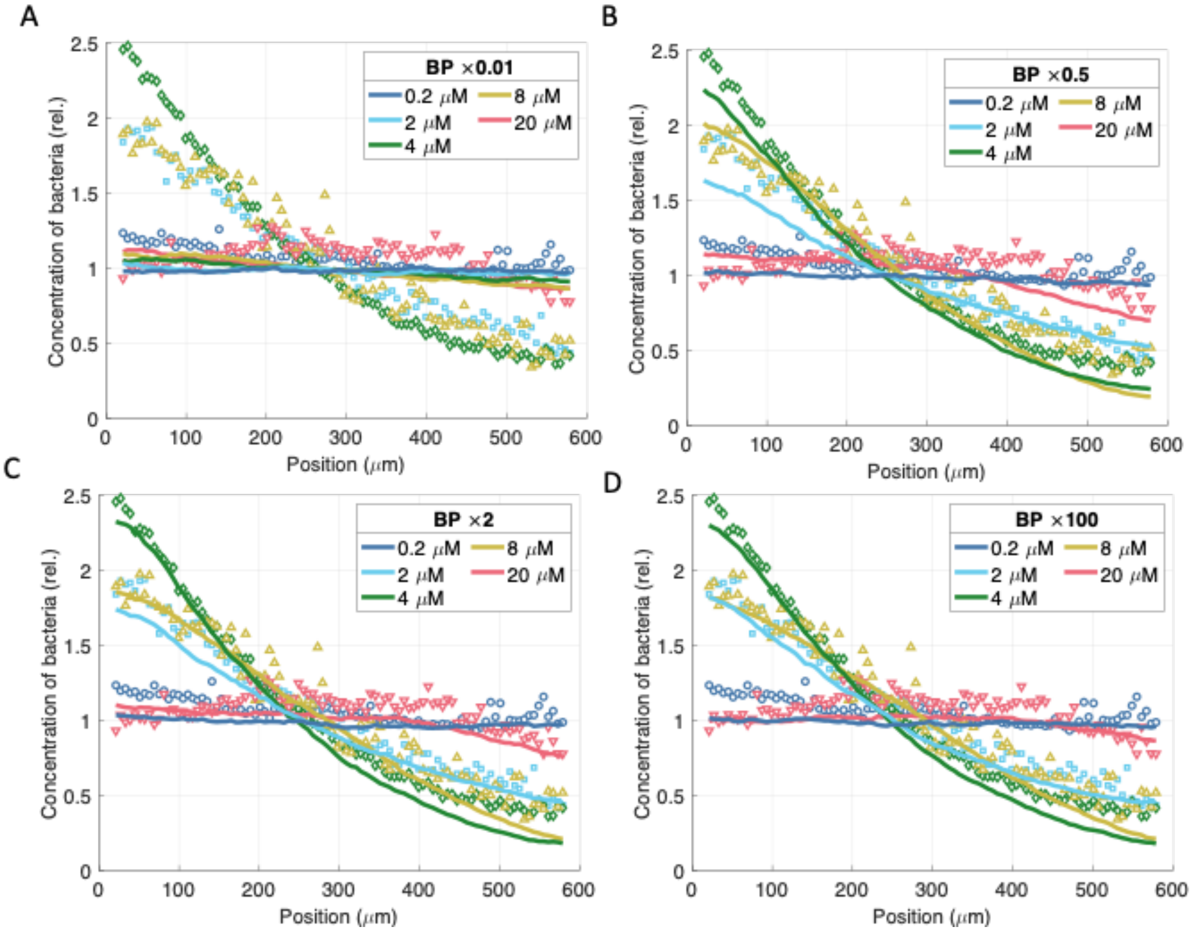


**Figure S6: Insensitivity of chemotactic response to binding protein abundance.** Although the chemotactic response disappears when the binding protein abundance, [BP], is drastically reduced by a factor of 100 (A), the chemotactic response is insensitive to both smaller variations in abundance (B) and increases in binding protein abundance (C&D). This insensitivity to binding protein abundance can be easily seen from the linear approximation of the chemotactic signal, [L:BP], as a function of $[L]_{ext}$: it is independent of [BP].

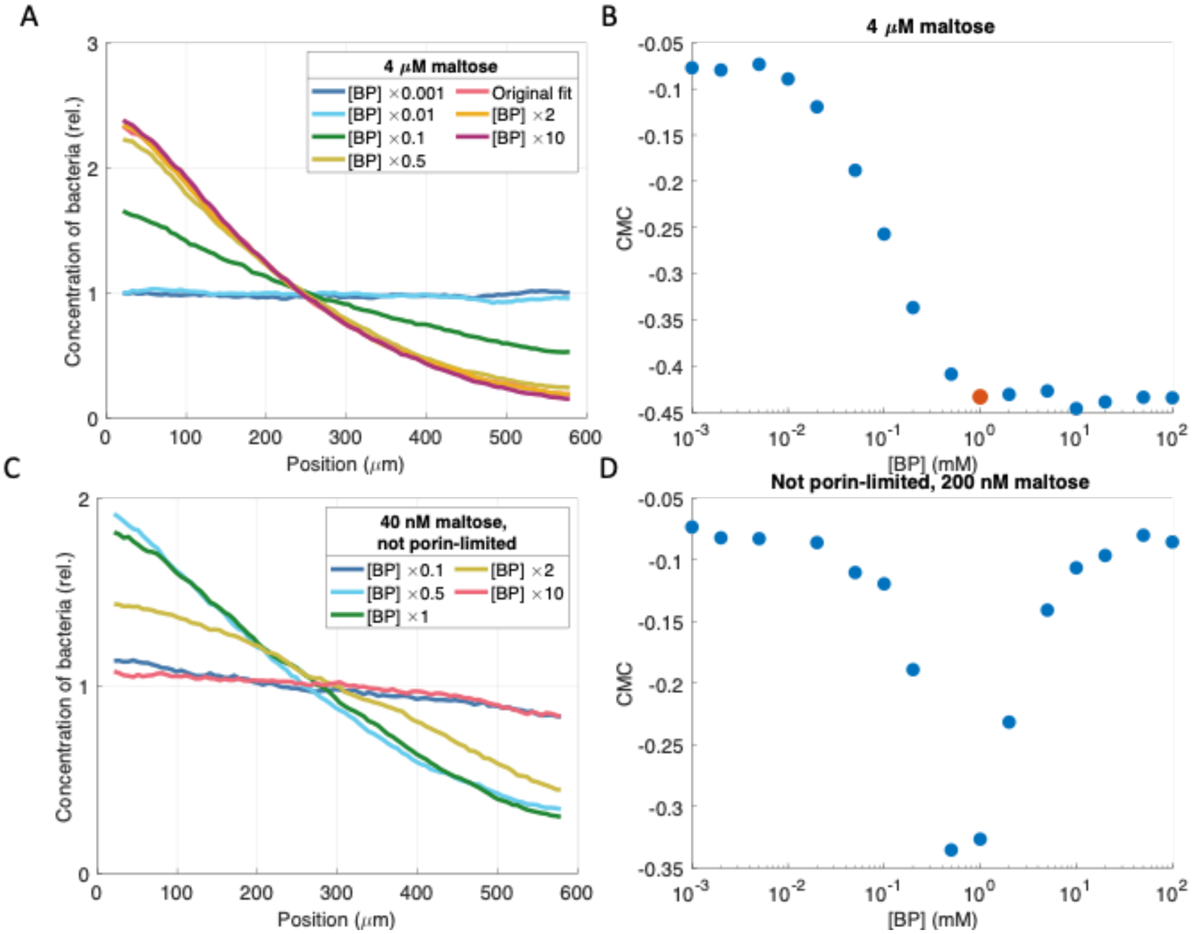


**Figure S7**: **Sensitivity of response to binding protein abundance.** Here we use our fitted SPECS model to predict the peak chemotactic response as a function of binding protein abundance. To quantify the response, we use the chemotaxis migration coefficient (CMC), which is defined as $\mathrm{CMC} = \frac{\langle x \rangle - 300}{300}$, where $\langle x \rangle$ is the average position in microns of the cells across the 600 μm channel. (A&B) Our model suggests that the CMC does not vary for sufficiently high binding protein abundances. The red dot indicates our estimate of the cells' average binding protein abundance and corresponding response at 4 μM maltose. (C&D) On the other hand, if instead transport were not porin-limited, the response would be highly sensitive to binding protein abundance.

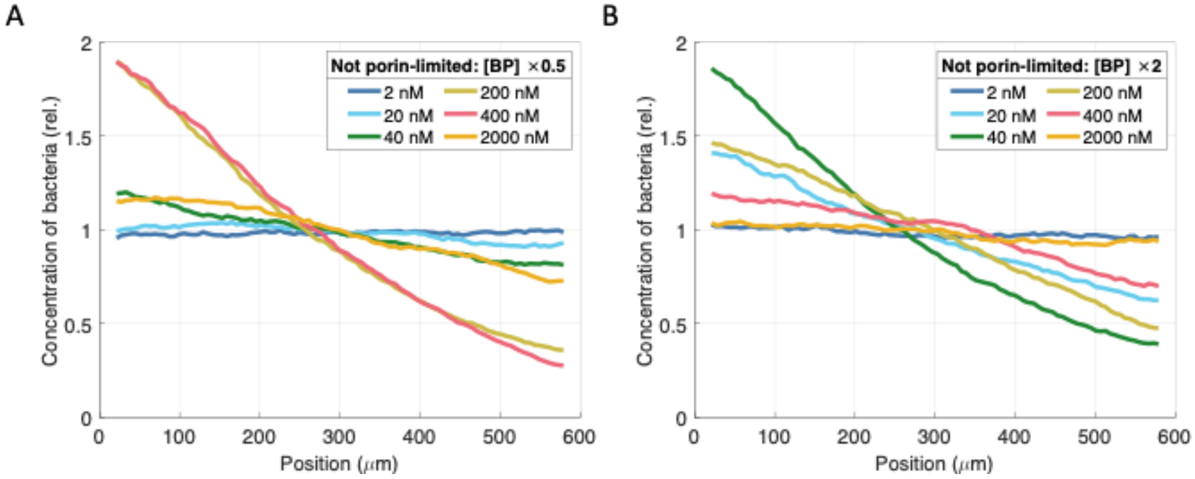


**Figure S8: Sensitivity of the chemotactic response to binding protein abundance when transport is not porin-limited.** If instead transport were not porin-limited so that $[L]_p \approx [L]_{ext}$, then the chemotactic response would be proportional to binding protein abundance and thus the response would be highly sensitive to variations in binding protein abundance.

**A** **B**

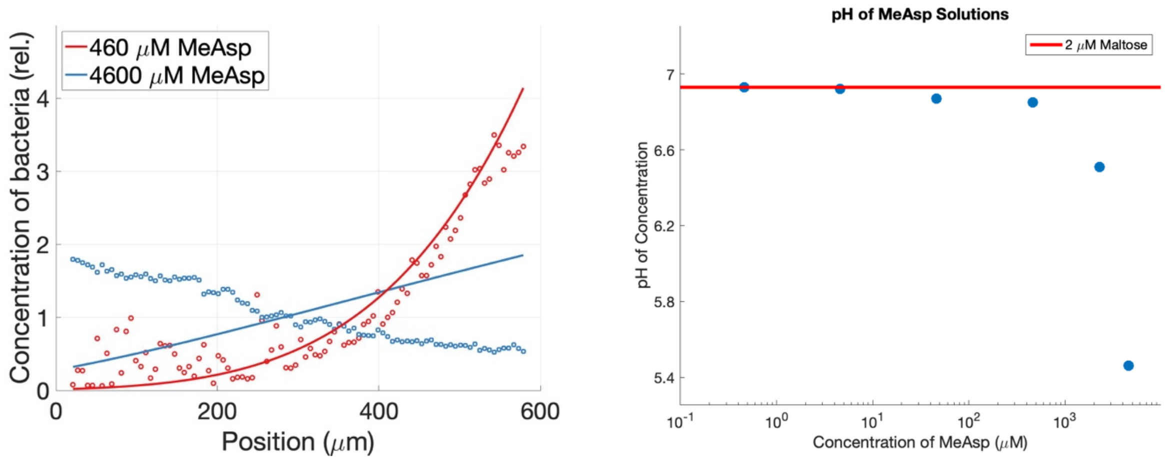


**Figure S9: Cells are repelled from high concentrations of MeAsp.** (A) Steady-state distributions from experimental chemotaxis assays in gradients of MeAsp along with predicted best-fit using analytical approximation with direct-binding model. (B) pH of MeAsp solutions used in experiments. Because we suspect pH taxis causes repulsion (Hu & Tu, 2014), we restricted our model fitting to concentrations of MeAsp less than 500 μM.

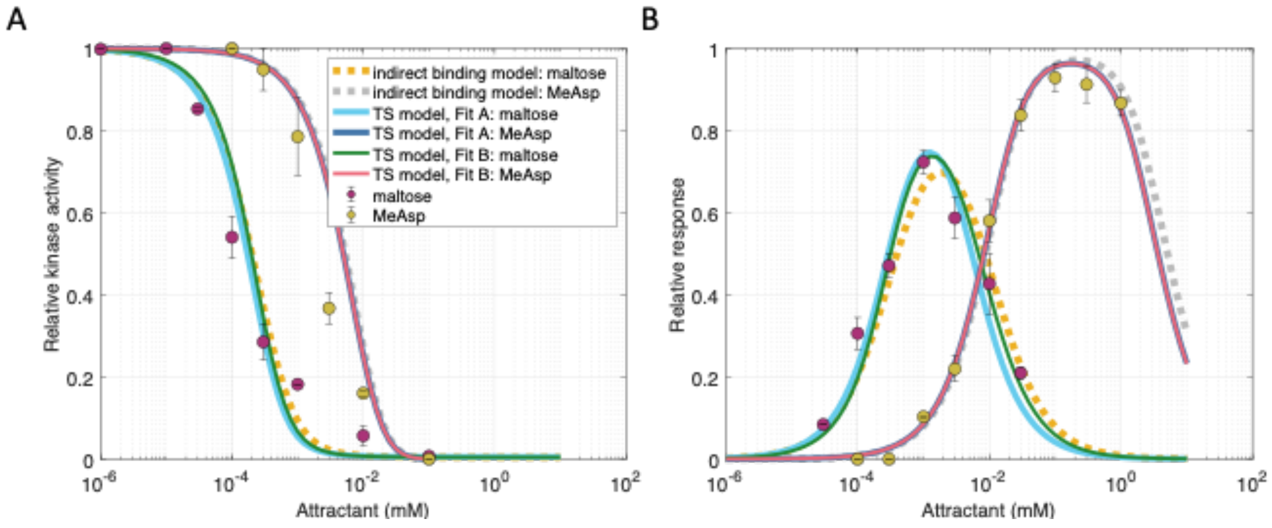


**Figure S10: Fitting the transport-and-sensing model to FRET activity assays.** Here we compare the best fits obtained by Neumann *et al.* using their indirect binding model and best fits we obtained using our transport-and-sensing (TS) model (Methods). The FRET assay data in (A) shows the dose response of *E. coli* LJ110 Δ(*cheY cheZ*) to step additions of maltose or methyl-aspartate (MeAsp), and (B) shows the dynamic range to three-fold step additions, during which the cells were adapted prior to each new addition (Neumann et al., 2010). The indirect-binding model assumes that the periplasmic concentration of free maltose equals the extracellular concentration, and its fit is with receptor cooperativity $n_{Tar} = 6$, binding protein dissociation constant $K_{BP} = 2$ μM, dissociation constant to binding protein ratios $K_{I,Mal}/[BP] = 0.4$ and $K_{A,Mal}/[BP] = 6$, and methyl-aspartate dissociation constants $K_{I,MeAsp} = 30$ μM and $K_{A,MeAsp} = 500$ μM. Fit A of our transport-and-sensing (TS) model uses parameters: $n_{Tar} = 6$, $K_{I,MeAsp} = 27.5$ μM, $K_{A,MeAsp} = 365$ μM, $K_{I,Mal} = 14.4$ μM, $K_{A,Mal} = 49.7$ μM, $[R]_{total} = 12.6$ μM, $[BP]_{total} = 101$ μM, and $V_c/V_p = 1.09 \times 10^{-5}$. Fit B uses parameters: $n_{Tar} = 6$, $K_{I,MeAsp} = 27.5$ μM, $K_{A,MeAsp} = 363$ μM, $K_{I,Mal} = 394$ μM, $K_{A,Mal} = 2{,}040$ μM, $[R]_{total} = 21.0$ μM, $[BP]_{total} = 1290$ μM, and $V_c/V_p$ = 1.00 × $10^{-5}$. Fit A correctly predicts maltose-Tar dissociation constants in the micromolar range but predicts low binding protein abundances. On the other hand, Fit B predicts reasonable protein abundances but much too high dissociation constants.

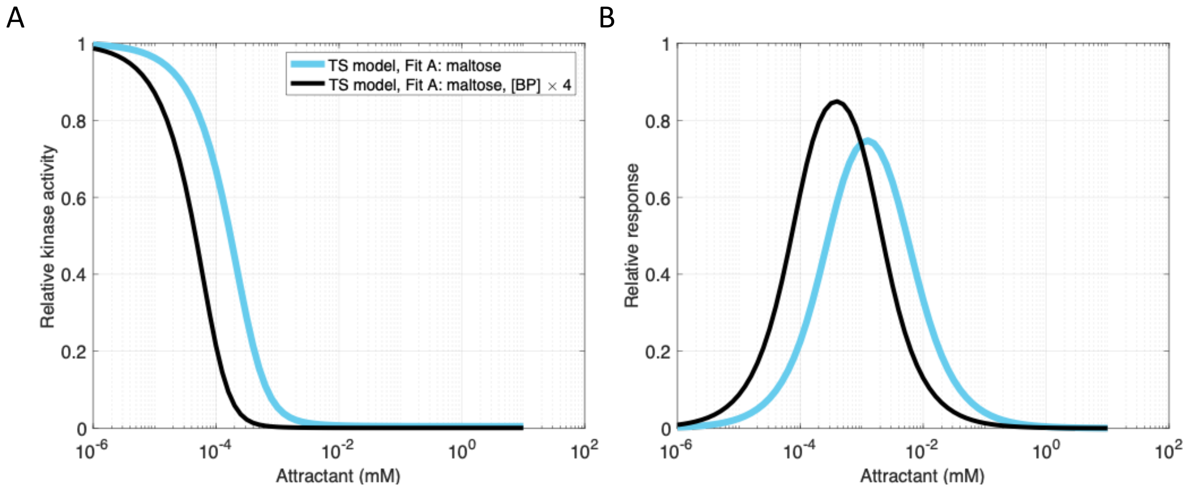


**Figure S11: Modifying binding-protein abundance for Fit A of transport-and-sensing model to FRET data.** Fit A of the transport-and-sensing model (Figure S10) predicts a binding-protein abundance that is a factor of ten lower than previous literature estimates. However, the finding from the FRET assays that increased binding-protein expression increases chemotactic sensitivity supports our hypothesis that binding-protein abundance was low for the strain and culture conditions of the FRET assays: our model predicts that, in this regime of low binding-protein abundance, chemotactic sensitivity increases with binding protein abundance.

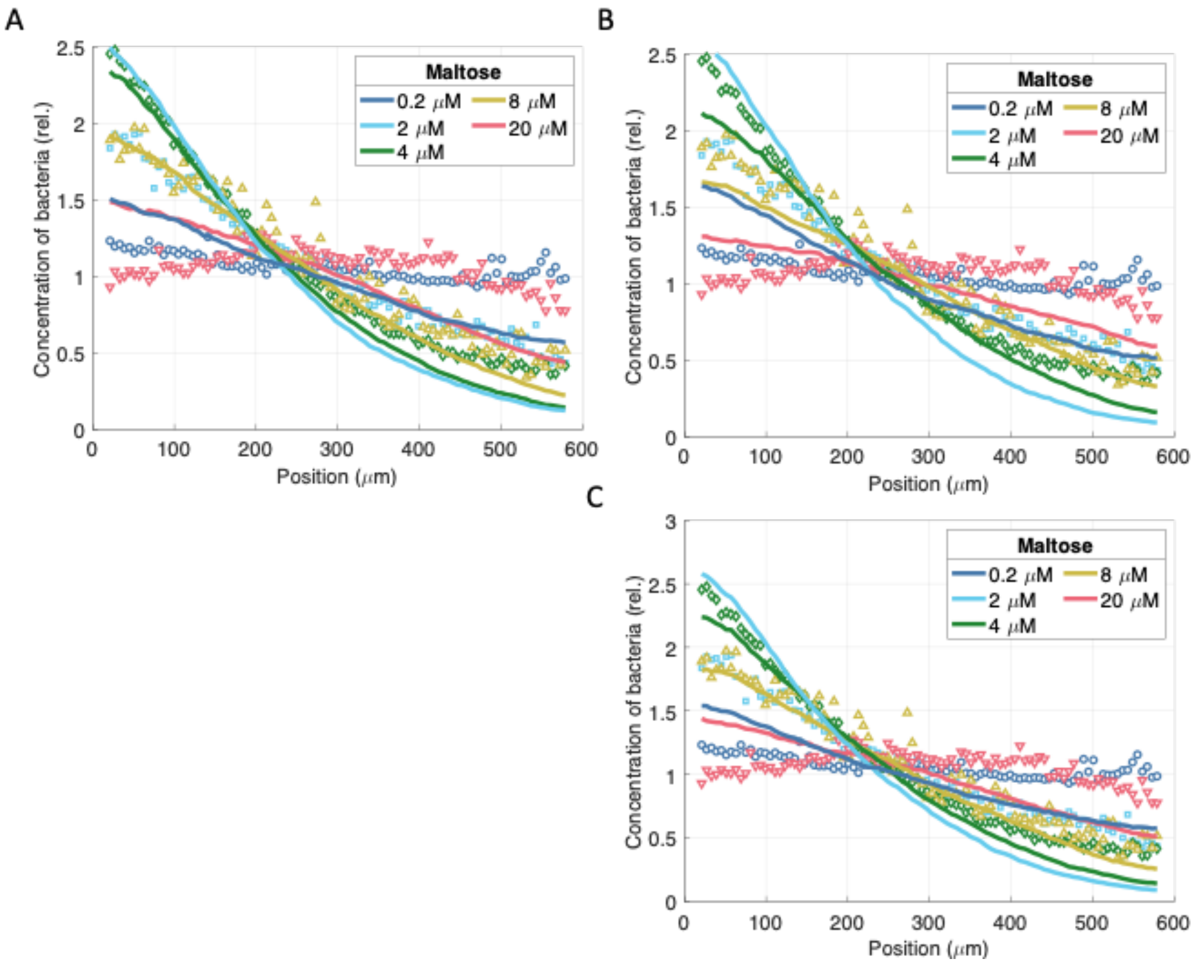


**Figure S12: Predicted steady-state distributions of cells in maltose gradients using SPECS simulator with fits using FRET activity level assays.** (A) Experimental cell distributions shown with predictions from SPECS simulator incorporated with indirect-binding model and using parameters from fitted FRET data: $n_{Tar} = 4$, $K_{BP} = 2$ μM, $K_{I,Mal}/[BP] = 0.4$, $K_{A,Mal}/[BP] = 6$, and $p_0 = 0.1$ (Neumann, et al. 2010). Same experimental cell distributions shown with predictions from SPECS simulator incorporated with transport-and-sensing model and using Fit A (B) or Fit B (C) obtained from fitting FRET data (Figure S10) with $n_{Tar} = 4$.
